# The structure of a calsequestrin filament reveals mechanisms of familial arrhythmia

**DOI:** 10.1101/672303

**Authors:** Erron W. Titus, Frederick H. Deiter, Chenxu Shi, Julianne Wojciak, Melvin Scheinman, Natalia Jura, Rahul C. Deo

**Author notes:** Correspondence (E.W.T.); (R.C.D.).

## Abstract

Mutations in the calcium-binding protein calsequestrin cause a highly lethal familial arrhythmia, catecholaminergic polymorphic ventricular tachycardia (CPVT). In vivo, calsequestrin multimerizes into filaments, but a compelling atomic-resolution structure of a calsequestrin filament is lacking. We report a crystal structure of a cardiac calsequestrin filament with supporting mutation analysis provided by an *in vitro* filamentation assay. We also report and characterize a novel disease-associated calsequestrin mutation, S173I, which localizes to the filament-forming interface. In addition, we show that a previously reported dominant disease mutation, K180R, maps to the same multimerization surface. Both mutations disrupt filamentation, suggesting that dominant disease arises from defects in multimer formation. A ytterbium-derivatized structure pinpoints multiple credible calcium sites at filament-forming interfaces, explaining the atomic basis of calsequestrin filamentation in the presence of calcium. This work advances our understanding of calsequestrin biochemistry and provides a unifying structure-function molecular mechanism by which dominant-acting calsequestrin mutations provoke lethal arrhythmias.

---

The Ca^2+^ ion is a ubiquitous chemical messenger in eukaryotic cells, where transient changes in intracellular calcium trigger a diverse set of signal transduction pathways. Facilitating this signaling role, the calcium gradient across the plasma membrane dwarfs the gradients of the other common ions, ranging over 5 orders of magnitude from approximately 2mM extracellularly to 100 nM intracellularly ***Carafoli and Krebs (2016)***. Many critical signaling pathways involve calcium efflux from the endoplasmic reticulum (ER), an organelle that maintains an ionic milieu resembling extracellular conditions. Direct physical coupling between the ER lumen and the extracellular space allows some degree of passive equilibration between these two compartments ***Prakriya and Lewis (2015)***, so that the biochemical work of establishing and maintaining calcium stores is not done twice.

In muscle, the transduction of an electrical signal at the membrane into a calcium flow that activates the contractile apparatus is known as excitation-contraction coupling. Under conditions of high muscle loading, calcium release from the sarcoplasmic reticulum (SR, a muscle-specific extension of the ER) becomes much more substantial than other biological calcium fluxes. In order to minimize the energy expended per contractile cycle, muscle cells possess enhanced calcium buffering at the sites of storage and release, allowing a high total calcium content with much lower free calcium. Calsequestrin (CSQ) is a 44 kD highly acidic protein that serves as the principal SR calcium buffer of cardiac and skeletal muscle, storing up to 50% of SR calcium in a bound state, with each calsequestrin monomer storing up to 40-50 calcium ions MacLennan and Wong (1971); ***MacLennan (1974)***; ***Ostwald and MacLennan (1974)***; ***Costello et al. (1986)***; ***Franzini-Armstrong et al. (1987)***; ***Wang et al. (1998)***; ***Park et al. (2004)***; ***Knollmann et al. (2006)***. Calsequestrin is complexed to the SR calcium channel, the ryanodine receptor (RyR), thereby ensuring that calcium is stored at the site of its release ***Bers (2004)***.

Since their initial identification, in 1971, a substantial field of research has formed around calsequestrins. Calsequestrins (CASQ1 in skeletal muscle and CASQ2 in cardiac muscle) are highly homologous in structure and function, and 64% identical in sequence, with skeletal muscle calsequestrin appearing to have higher calcium capacity ***Park et al. (2004)***. Calcium-binding propensity is explained in large part by calsequestrin’s remarkable fraction of negatively-charged acidic residues (26% glutamate or aspartate in CASQ2 and 28% in CASQ1, with corresponding average isoelectric points of 4.2 and 4.0, respectively). In both cardiac and skeletal muscle, calsequestrin localizes to the junctional SR (jSR) of muscle and forms multimers that are anchored to the luminal SR membrane. From low resolution electron micrographs of isolated skeletal muscle fibers, these multimers appear as a dense collection of filaments ***Franzini-Armstrong et al. (1987)***; ***Perni et al. (2013)***. These filaments are anchored to a complex consisting of RyR and the single-pass transmembrane proteins triadin and junctin ***Bers (2004)***. Extended homo-multimerization provides high-density calcium storage, but multimerization appears to be essential for localization, too, in that multimerization-defective mutants are trafficked along the secretory pathway and lost from the ER/SR ***Milstein et al. (2009)***; ***McFarland et al. (2010)***; ***Knollmann (2010)***. Additional insights into calsequestrin biochemistry and cell biology encompass post-translational modifications ***Sanchez et al. (2012b)***; ***Kirchhefer et al. (2010),*** calcium-storage capacity ***Park et al. (2004),*** and interactions with the calcium release unit ***Zhang et al. (1997)***; ***Rani et al. (2016)***; ***Handhle et al. (2016)***.

Calsequestrin’s relevance to human disease is well established. Mutations in skeletal muscle calsequestrin have been putatively linked to malignant hyperthermia and to vacuolar myopathies ***Lewis et al. (2015),*** while mutations in cardiac calsequestrin are well known to cause catecholaminergic polymorphic ventricular tachycardia (CPVT), a highly lethal familial arrhythmia. CPVT is caused by a state of cardiac calsequestrin deficiency, whether arising from null/hypomorphic alleles or from point mutants that disrupt calsequestrin multimerization ***Bal et al. (2010)***; ***Bal et al. (2011)***. Consistent with a mechanism where by deficiency leads to disease, most calsequestrin-associated CPVT mutations have recessive inheritance. For a protein whose function depends on multimerization, there is, at least to date, a surprising paucity of mutations with putative dominant negative mechanism.

Despite the rich body of work concerning calsequestrin biology, no compelling high-resolution candidate structure for a calsequestrin filament has emerged. Although sixteen crystal structures of calsequestrins have been published ***Wang et al. (1998)***; ***Park et al. (2004)***; ***Kim et al. (2007)***; ***Sanchez et al. (2012b)***; ***Sanchez et al. (2012a)***; ***Lewis et al. (2015)***; ***Lewis et al. (2016)***, none of these contain a convincing filament-like assembly. All 16 prior structures reveal calsequestrin dimers that are nearly-identical to one another, but a search for dimer-to-dimer interfaces within and across these crystal unit cells reveals only weak crystallographic packing contacts that appear incompatible with robust biological multimerization. In addition, the observed dimer-to-dimer interfaces vary substantially from one lattice to another. Critically, a lack of mutagenesis studies supporting proposed oligomerization interfaces in the prior published structures calls into question whether the relevant biological multimer has ever been observed. Cumulatively, the prior studies have established that calsequestrins dimerize in a wide variety of conditions, with ***intra-dimer*** interactions that are largely the same across published structures, but the mechanism by which dimers assemble into higher order multimers (***inter***-dimer assembly) remains elusive.

A structure of a calsequestrin filament would advance our understanding of calsequestrin biology and permit a comprehensive mapping of disease-causing mutations to biologically-relevant multimerization interfaces. We report here an investigation of dominant-acting cardiac calsequestrin disease mutants that has culminated in a new X-ray structure of cardiac calsequestrin, one that we believe reveals the biologically relevant filament-forming interface. We show that known dominant-acting mutants - one previously reported, and one described for the first time in this study - map to the newly reported multimerization interface. Furthermore, we provide supporting biochemical analysis of likely calcium-binding sites where the protein multimerizes. These findings fundamentally advance our understanding of the mechanism by which calsequestrin contributes to calcium homeostasis and provide insight into mechanisms of calsequestrin-associated familial diseases.

## Autosomal Dominant *CASQ2* Disease Mutations Disrupt Calsequestrin Multimerization Yet Are Not Well-Explained by Prior Calsequestrin Structures

We encountered a proband from a family with CPVT-like tachy-arrhythmias and multiple cases of sudden, unexplained death at a young age (Figure 1a). The proband presented at age 33 with a biventricular tachycardia on ECG. She underwent an electrophysiologic study that revealed an inducible ventricular tachycardia with focal origination next to the anterior fascicle, and an inducible atypical atrioventricular nodal reentry tachycardia that was successfully ablated. Targeted sequencing of channelopathy genes *(KCNQ1, KCNH2, SCN5A, ANK2, KCNE1, KCNE2, KNCJ2, CAV3, RYR2,* and *CASQ2*) revealed only a heterozygously-carried isoleucine-for-serine substitution at position 173 in cardiac calsequestrin. The proband’s son and multiple other family members report a tachycardic phenotype (ranging from self-reported palpitations to diagnosed episodes of tachycardia). The family history is also notable for multiple cases of sudden cardiac death at a young age. Overall, the distribution of affected individuals in the pedigree is potentially consistent with dominant inheritance. As the family did not consent to follow-up genetic testing or clinical phenotyping, we were unable to rigorously assess disease/mutation co-segregation.

**Figure 1:**
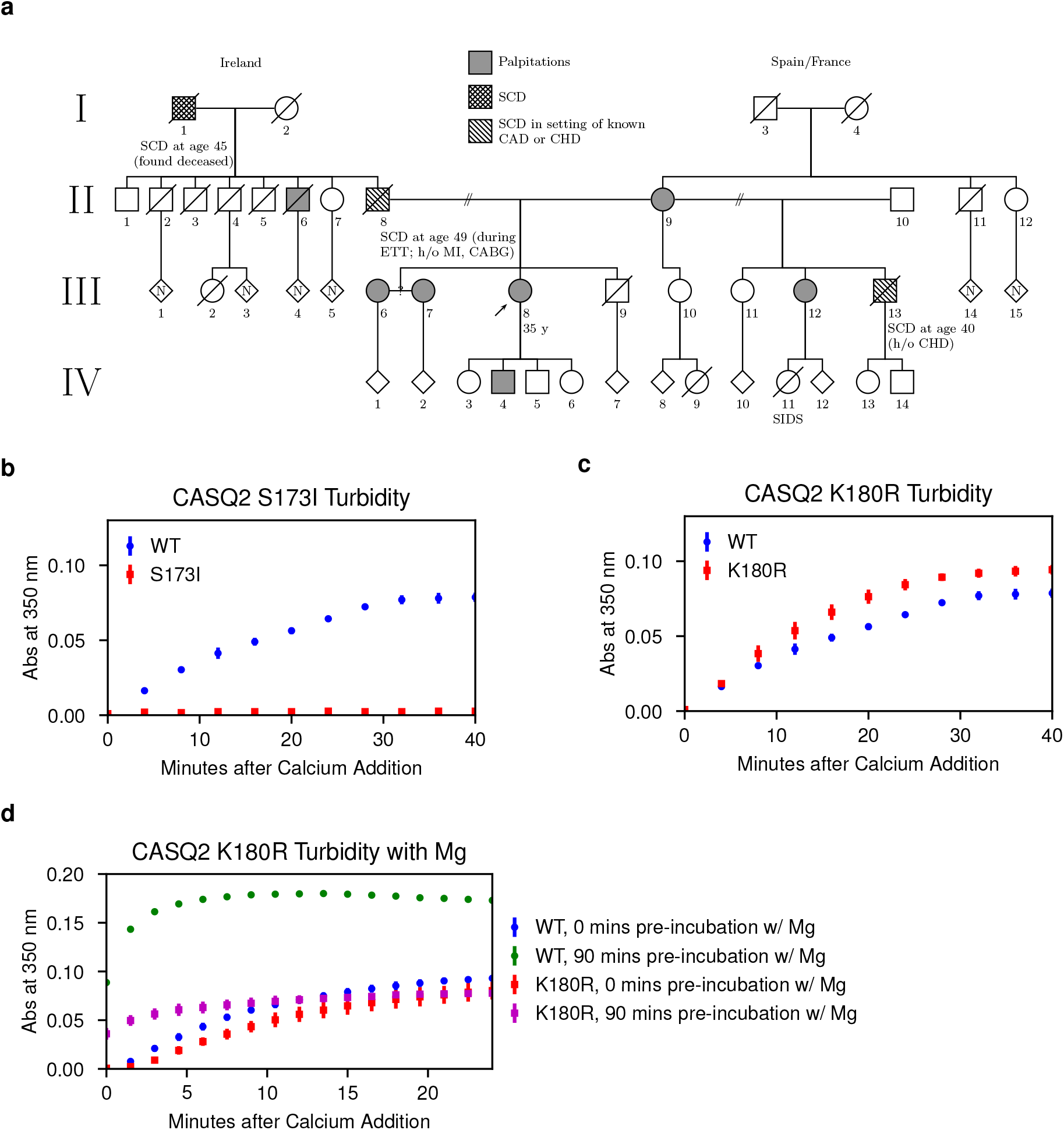
Autosomal Dominant *CASQ2* Disease Mutations Disrupt Calsequestrin Multimerization Yet Are Not Well-Explained by Prior Calsequestrin Structures. **a**, Pedigree of a large extended family with the S173I mutation and a CPVT-like phenotype. CABG = coronary artery bypass graft; CAD = coronary artery disease; CHD = congenital heart disease; ETT = exercise treadmill test; MI = myocardial infarction; SCD = sudden cardiac death; SIDS = sudden infant death syndrome. **b**, Multimerization kinetics of the S173I mutant observed using a turbidity assay after addition of 1 mM CaCl_2_ to purified protein (pH 7.4, 20 mM NaCl, 85 mM KCl). c, Multimerization kinetics of the K180R mutant observed using a turbidity assay (same conditions as in **b**). **d**, Multimerization kinetics of the K180R mutant observed using a turbidity assay (same conditions as in B, but with 2mM MgCl_2_ added prior to calcium).

The genetic evidence for pathogenicity of the S173I novel variant is not conclusive, but since known pathogenic point mutations in *CASQ2* exhibit defective filamentation, we elected to investigate the S173I mutation biochemically in a simple turbidity assay. The turbidity assay for S173I reveals a profound decrease in multimerization rate (Figure 1b). Minimal improvement in the multimerization rate is observed in a non-physiologic 0 mM potassium condition (Figure S1), demonstrating that the protein is intact but defective in multimerization.

The sole known *CASQ2* disease mutation with strong evidence for dominant inheritance, K180R, was not reported to cause defects in multimerization and was proposed to act via alternative mechanisms *Gray et al. (2016).* The striking effect of S173I on CASQ2 mutimerization, combined with the fact that other known *CASQ2* point mutants exhibit the same biochemical defect, prompted us to reexamine multimerization capacity of the K180R mutant. The turbidity assay for K180R under the same conditions used for S173I shows little difference from the wild type *CASQ2* variant (Figure 1c). However, prior reports suggested that calsequestrin maintains distinct magnesium and calcium binding sites ***Krause et al. (1991)***. Therefore, we investigated filamentation kinetics of the K180R mutant in the presence of magnesium. Strikingly, extended incubation of the K180R mutant with magnesium (2 mM MgCl2) prior to addition of calcium yields a profound filamentation defect (Figure 1d). In vivo, calsequestrin would likely encounter similar levels of free magnesium throughout the SR.

Interestingly, neither S173 nor K180 fall at credible, previously identified candidate multimerization interfaces: they are not near the intra-dimer interface, nor are they near candidate inter-dimer interfaces in the prior crystal structures. As the dominant inheritance pattern is classically associated with disease mutations that disrupt protein-protein interactions, we would have expected to find these and other dominant-acting disease mutations at cardiac calsequestrin’s dimer or multimer interfaces. We therefore elected to pursue another structure in the belief that the biologically relevant multimer has not yet been observed.

## The New Cardiac Calsequestrin Filament Candidate Is Helical at the Domain Level

We have determined a new crystal structure of human cardiac calsequestrin obtained from a full-length construct in a very low-pH (3-3.5) crystallization condition. The previously characterized calsequestrin dimer is again observed, but now in an arrangement that produces a closely-packed filament (Figure 2a). Crystallographic data collection and processing statistics are summarized in (Table S1) for the native structure as well as a ytterbium-soaked condition used to identify probable calcium-binding sites.

**Figure 2:**
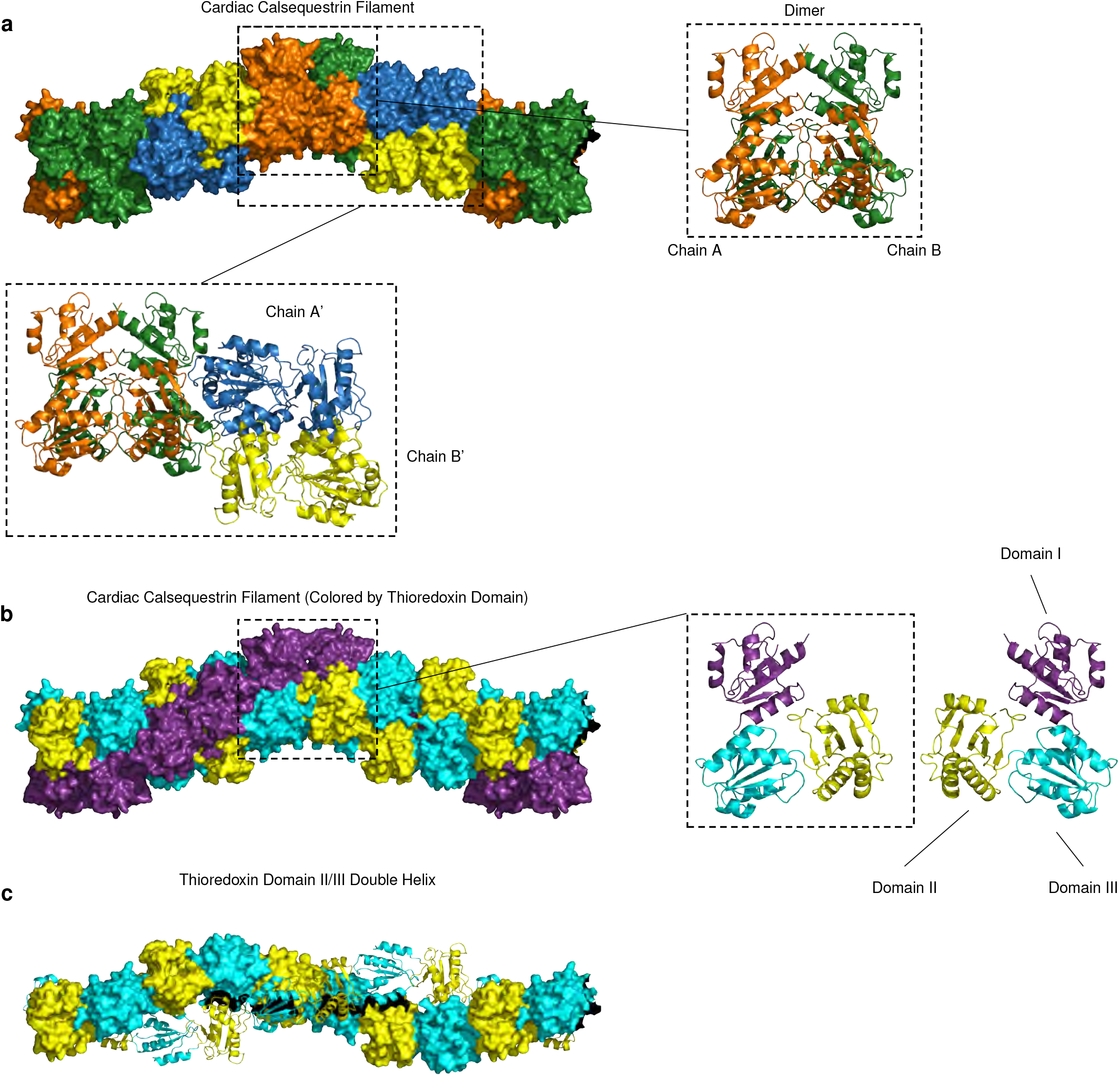
The New Cardiac Calsequestrin Filament Candidate Is Helical at the Domain Level. **a**, The cardiac calsequestrin candidate filament (PDB ID 6OWV) is shown, along with a representative dimeric and tetrameric assembly. Dimers are stacked on a screw axis with 90 degrees of rotation per dimer. **b**, The helical character of the filament is revealed at the domain level. Cardiac calsequestrin monomers are colored by thioredoxin domain (domain I, purple; domain I, cyan; domain III, yellow). Viewed at the level of its thioredoxin domains (3 per protomer), the filament consists of an inner thioredoxin double helix (domains II and III) with an outer thioredoxin single helix (domain I) wrapping the double helical core. Right side: The monomers are translated but remain in their dimer-forming orientation. **c**, The inner double helix of the filament consisting of thioredoxin domains II and III.

The new structure provides a compelling candidate for a biologically-plausible higher-order multimer. The repeating unit of the native crystal resides in a higher-symmetry point group compared to prior calsequestrin structures (Table S2). The oligomer-forming contacts that exist between dimers are novel, differing significantly from all previously reported calsequestrin crystal structures (Figure S2). Furthermore, the candidate filament-forming interfaces in our structure collectively encompass significantly greater buried surface area than observed at any previous calsequestrin inter-dimer interface (Figure S2).

The cardiac calsequestrin monomer, like its skeletal calsequestrin equivalent, consists of an N terminal loop, 3 thioredoxin domains, and a disordered acidic tail. Within the dimer, there is two-fold symmetry with mutual exchange of N-terminal loops, as previously observed (Figure 2b). In our structure, the dimers are stacked along a screw axis to form the filament, with each dimer rotated 90° with respect to its neighbors. Although the dimers are positioned at discrete 90° rotations with respect to one another, the underlying architecture of the filament is in fact helical at the level of thioredoxin domains (Figure 2b). Thioredoxin domains II and III form a double helix at the core of the filament (Figure 2c). An outer helix or “collar” of larger diameter, consisting only of thioredoxin domain I, then winds around the inner double helix. All helices are left-handed, corresponding to the left-handed screw axis at the level of dimer-stacking.

## The 3-Helix Configuration of the New Filament Candidate Promotes Close-Packing of Thioredoxin Domains

The 3-helix configuration appears to promote the close packing of globular thioredoxin domains. Helical packing permits each domain to contact multiple other domains in multiple other protomers, which is in stark contrast to other reported candidate structures (PDB ID 1A8Y, rabbit skeletal muscle calsequestrin; PDB ID 1SJI, canine cardiac calsequestrin), which lack the helical pitch and have much more limited interacting surface area (Figure 3). The close-packing of the new filament candidate is starkly visible when thioredoxin domains are represented as equally-sized spheres centered at the domain center of mass (Figure 3, right-hand side). Prior putative calsequestrin filament candidates have fewer interdomain contacts, fewer inter-protomer contacts, and substantially less buried surface area compared to the new candidate.

**Figure 3:**
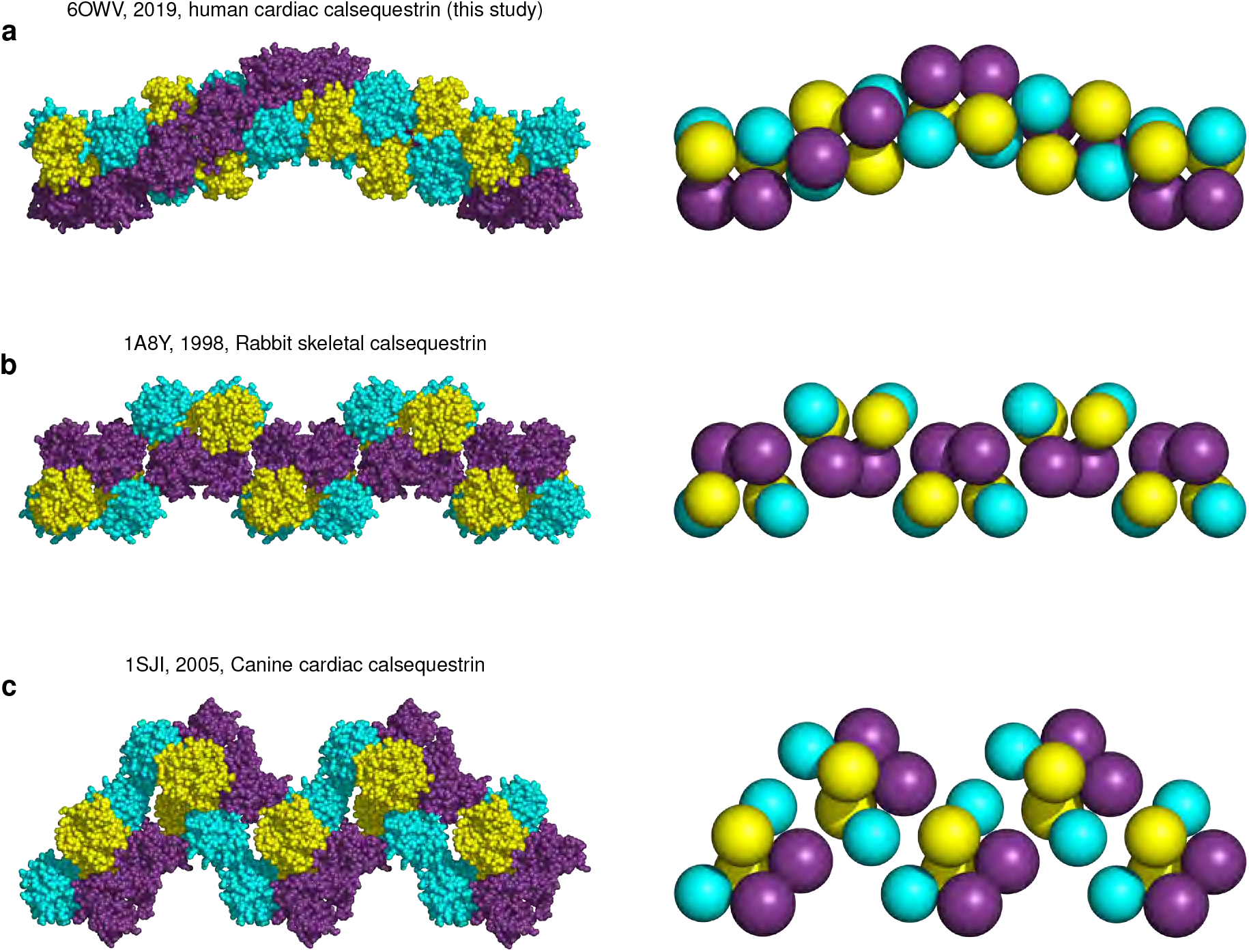
The 3-Helix Configuration of the New Filament Candidate Promotes Close-Packing of Thioredoxin Domains. **a**, The new candidate cardiac calsequestrin filament assembled from crystallographic symmetry operations on 6OWV (human CASQ2, 2019). The new candidate CASQ2 filament exhibits tight packing of protomers and thioredoxin domains (shown on the right using equal-size spheres placed at the center of mass of each thioredoxin domain). **b**, A putative skeletal calsequestrin filament assembled from crystallographic symmetry operations on 1A8Y (rabbit CASQ1, 1998). Right-side: equal-size spheres represent thioredoxin domains. **c**, A putative skeletal calsequestrin filament assembled from crystallographic symmetry operations on 1SJI (canine CASQ2, 2005). Right-side: equal-size spheres represent thioredoxin domains.

Notably, our crystallization condition produced crystals only at a low pH (3-3.5, measured by applying a crystallization drop to litmus paper) resulting from the use of aged PEG reagents (presumably having undergone degradation to glycolic acid) or from direct addition of concentrated HCl. This restriction obtained irrespective of calcium in concentrations ranging from trace to 14 mM (higher calcium concentrations led to precipitation, likely due to the presence of sulfate, and were therefore not assessed). Given calsequestrin’s remarkable overall acidity and low isoelectric point, an explanation for the role of low pH in promoting calsequestrin filamentation could be the neutralization of substantial negative charge on acidic side chains that extend across the filament interfaces.

## Lanthanide Substitution Reveals Cation-Binding Sites Within Filamented Calsequestrin Dimers

Prior work has identified putative calcium binding sites at the dimer interface of calsequestrin ***Sanchez et al. (2012a)***. Our new filament structure permits us to examine the role of multivalent ions at all surfaces relevant for filamentation - both the intra-dimer surfaces as well as the inter-dimer surfaces responsible for higher-order multimers. To localize candidate calcium ligand sites within the context of the new filament structure, we collected data from a Ytterbium (Yb)-soaked crystal (Table S1). From the anomalous map, we identified approximately 8 sites with strong Yb signal per CASQ2 chain (> 3.8 *σ* in the anomalous map and difference density in the Fo-Fc map). As the prior work to identify calsequestrin’s calcium-binding sites was based largely on the indirect method of inference from metal coordination geometry with nearby side chains and waters, the use of Yb provides a direct approach to confirming the prior findings, as well as extending our understanding to the entire filament.

We first assessed the presence of Yb at the intra-dimer interface. We identified several high-occupancy Yb sites, most of which are clustered in a narrow region between protomers, where they are coordinated by multiple, highly-conserved acidic residues (Figure 4 with supporting electron density maps in Figure S3). Consistent with the location of a putative calcium ion in the prior study, we find a Yb atom coordinated primarily by E147 of chain A and D278 of chain B. In addition, we find a Yb atom coordinated primarily by E143 and E275, as well as another Yb atom coordinated by D310 and a putative sulfate anion within a solvent cavity enclosed by the dimer. These Yb-bridging sites stabilize the interaction between thioredoxin domain II from protomer A and thioredoxin domain III of protomer B. The interactions between domains result in close juxtaposition of acidic side chains (Figure 4a), requiring neutralization of the charge by the multivalent counterions or, in the case of the native structure, protonation. In the native structure, the E147/D278 interaction provides an example of a carboxyl-carboxylate bond stabilized by low pH ***Sawyer and James (1982);*** Krause et al. (1991). Anomalous signal is found at this site in the derivative structure, but consistently on just one side of the otherwise symmetric dimer. This subtle asymmetry, with a cation bound on one side and a carboxyl-carboxylate bond on the other, likely reflects a degree difference in occupancy rather than a sharp distinction, but it is a consistent feature of the density map across all 4 dimers in the crystal asymmetric unit and may be conformationally conducive to filament formation.

**Figure 4:**
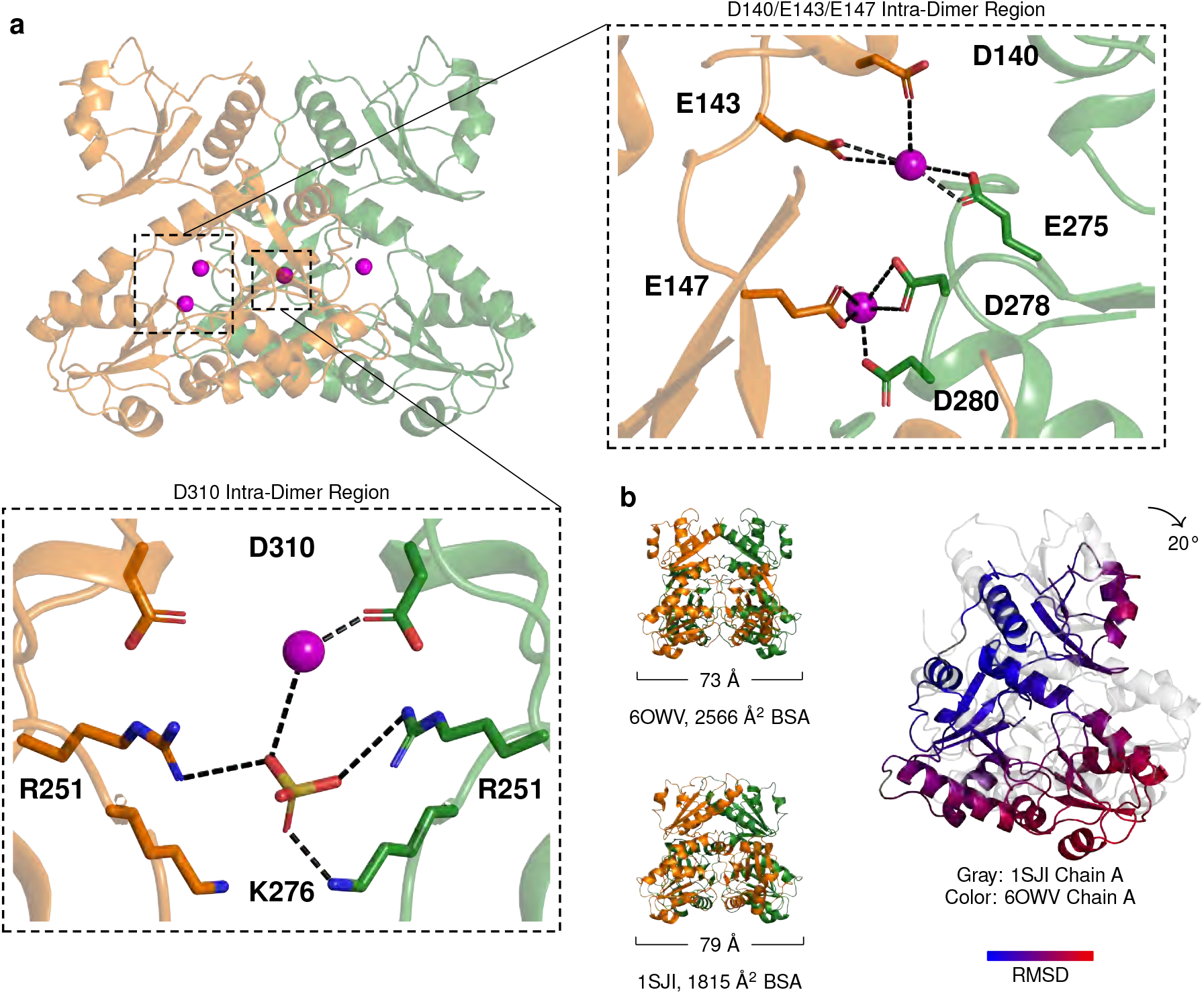
Lanthanide Substitution Reveals Cation-Binding Sites Within Filamented Calsequestrin Dimers. **a**, Dimer with ytterbium (Yb) sites (magenta spheres) within its interior cavity. Closeups focus on Yb positions that bridge dimer chains A and B. **b**, Comparison of a previously published cardiac calsequestrin dimer (1SJI) to the more tightly-packed dimer that we report. The tightly-packed dimer results primarily from rigid body rotation of the dimer chains inward (for a single chain, we observe 20° counter-clockwise rotation in the plane of the page when the other chain is fixed to the reference dimer). The inward rotation produces an increase in buried surface area (BSA) in thioredoxin domains II and III.

In both the native structure and Yb-soaked structure, the dimer subunits have undergone substantial inward rotation in comparison to the other high-resolution cardiac calsequestrin structure (PDB ID 1SJI) (Figure 4b). The inward domain movement is produced largely by rotation of the monomer as a rigid body and results in a conformation where domains II and III are more closely packed. This conformational shift recapitulates a similar finding from the prior study where a 15° rotation within the skeletal calsequestrin dimer was observed for structures crystallized in a high-calcium buffer ***Sanchez et al. (2012a)***. Upon inspection of all prior published calsequestrin structures, it is apparent that 6 prior calsequestrin structures belong to this “tightly-packed dimer” group, while the remaining 10 contain dimers that are more loosely-packed, without inward rotation of chains (Figure S4). The more tightly-packed structures were all crystallized in the presence of multivalent cations (usually calcium), or in one case (PDB ID 2VAF) a monovalent cation at extremely high concentration (2 M NaCl). The other group were crystallized with no added multivalent cations, with the exception of PDB ID 3TRP, which contained calcium in the crystallization drop at approximately 5-fold lower than lowest concentration from the tightly-packed group. Within the tightly-packed group, there is a greater degree of conformational disorder in Domain I (Figure S5). This is accompanied by a modest loss of contact in Domain I, while multiple hydrophobic side chains from Domains II and III that were not buried now obtain buried surface area (Figure S6). Thus, our data provide additional evidence for conformational change within the dimer upon calcium-binding (likely induced by closer approximation of acidic side chains), independent confirmation of intra-dimer calcium binding sites, and an explanation for why an altered conformation of the dimer becomes energetically tolerable in the presence of calcium.

## Lanthanide Substitution Reveals Cations Trapped at Inter-Dimer Filament-Forming Interfaces

Like the dimer interface, the inter-dimer filament-forming interface is bridged by Yb sites surrounded by closely-apposed acidic side chains (Figure 5 with supporting electron density maps in Figure S7). In contrast to the dimer interface, where carboxyl-carboxylate bonds are still seen in the Yb-derivative structure, the sites of carboxyl-carboxylate interaction at the inter-dimer interface have been thoroughly substituted with a cation. Identification of these coordination sites provides a testable model for the biological correctness of the new filament structure. To test this model, we mutated the Yb-binding aspartate or glutamate side chains to alanine and examined the effect of these mutations on filamentation kinetics. The most pronounced effects resulted from mutations targeting symmetrical interactions with Yb atoms within a large solvent cavity formed by the tetramer interface (Figure 5a). Specifically, residues D144 and E174 appear to coordinate Yb atoms within the cavity (the precise geometry of E174 is difficult to discern, due to lack of side chain density in the electron density maps of both the native and Yb-soaked structures). Mutation of these residues to alanine results in a dramatic increase in filamentation rate (Figure 5b). Residues E184 and E187 coordinate Yb atoms near the mouth of the opening, only partially shielded from exposure to the bulk solvent. Mutation of these residues results in a profound filamentation defect (Figure 5c). Glutamates 184 and 187 belong to an alpha helix that provides a linkage between domains of the outer thioredoxin collar. This helix, belonging to thioredoxin domain 2, sits between thioredoxin I domains of different dimers and interacts with inter-dimer salt bridges on either side. Alanine mutagenesis of the D50 residue that participates in a salt bridge with K180 at the N-terminal end of this helix produces a similar defect (Figure 5c, right-hand side).

**Figure 5:**
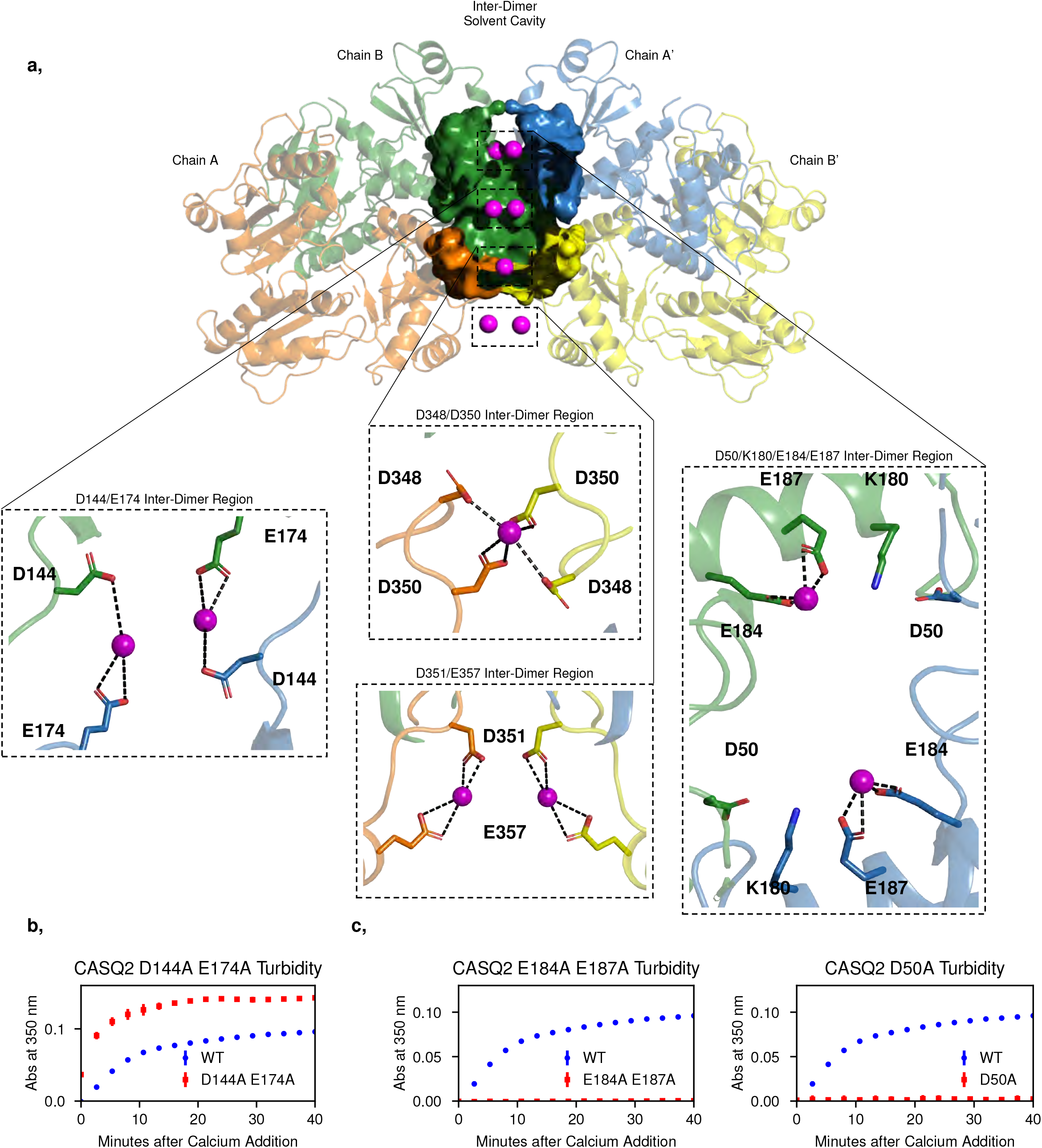
Lanthanide Substitution Reveals Cations Trapped at Inter-Dimer Filament-Forming Interfaces. **a**Yb (magenta spheres) bound within the walled pocket formed by the inter-dimer interface, with closeups of ytterbium site at E184 and E187, D348 and D350, and D351 and D357. Thioredoxin domain II of chain A’ (blue) is omitted to allow visualization of the interior of the solvent pocket formed by the inter-dimer interface. **b**Turbidity assay after alanine mutagenesis of putative calcium-binding residues D144 and E174. cLeft: turbidity assay after alanine mutagenesis of putative calcium-binding residues E184 and E187. Right: turbidity assay after alanine mutagenesis of the D50 residue that participates in a salt bridge with K180, adjacent to the E184/E187 ligand site.

Two more sites of strong Yb signal are notable. Residues D348 and D350 are oriented along with their symmetry mates to form a cluster of 4 acidic side chains that coordinate a single cation (Figure 5a and Figure S7). This Yb site forms the base of the large solvent cavity enclosed by the inter-dimer interface. Mutating the Yb-coordinating residues to alanine would be expected to relieve mutual repulsion of closely-packed acidic side chains. Consistent with this, alanine mutagenesis of these residues has a largely net neutral effect on filamentation kinetics (Figure S8a). Outside the inter-dimer cavity, on the fully solvent-exposed exterior of the filament, residues D351 and E357 and their symmetry mates form bidentate interactions with two Yb atoms, adopting conformations in which opposing acidic rotamers are bent away from one another, thereby alleviating electrostatic repulsion that would otherwise disrupt multimer formation (Figure 5a and Figure S7). As with the D348/D350 site, the bound cation at this site appears to facilitate filamentation by neutralizing the negatively charged surface of calsequestrin. Alanine mutagenesis of the D351 and E357 residues also has a largely net neutral effect on filamentation kinetics (Figure S8b).

## The Cardiac Calsequestrin Filament Possesses a Continuous, Solvent-Accessible Interior Cavity That Winds Along Its Long Axis

Intriguingly, the majority of Yb sites that we identify are located within a continuous solvent cavity that winds through the interior of the filament, forming a hollow core with solvent exposure at the intra-dimer and inter-dimer interfaces. Each dimer contains a solvent pocket within its interior, while another solvent pocket is formed at inter-dimer interface (Figure 6a-b). The stacking of dimers extends the cavity along the entire length of the filament (Figure 6c). The calcium ions that would appear to be bound in the dimer’s interior would constitute a separate store of calcium that is more slowly-mobilized than the highly-accessible pool of ions bound to the surface and the solvated acidic tail.

**Figure 6:**
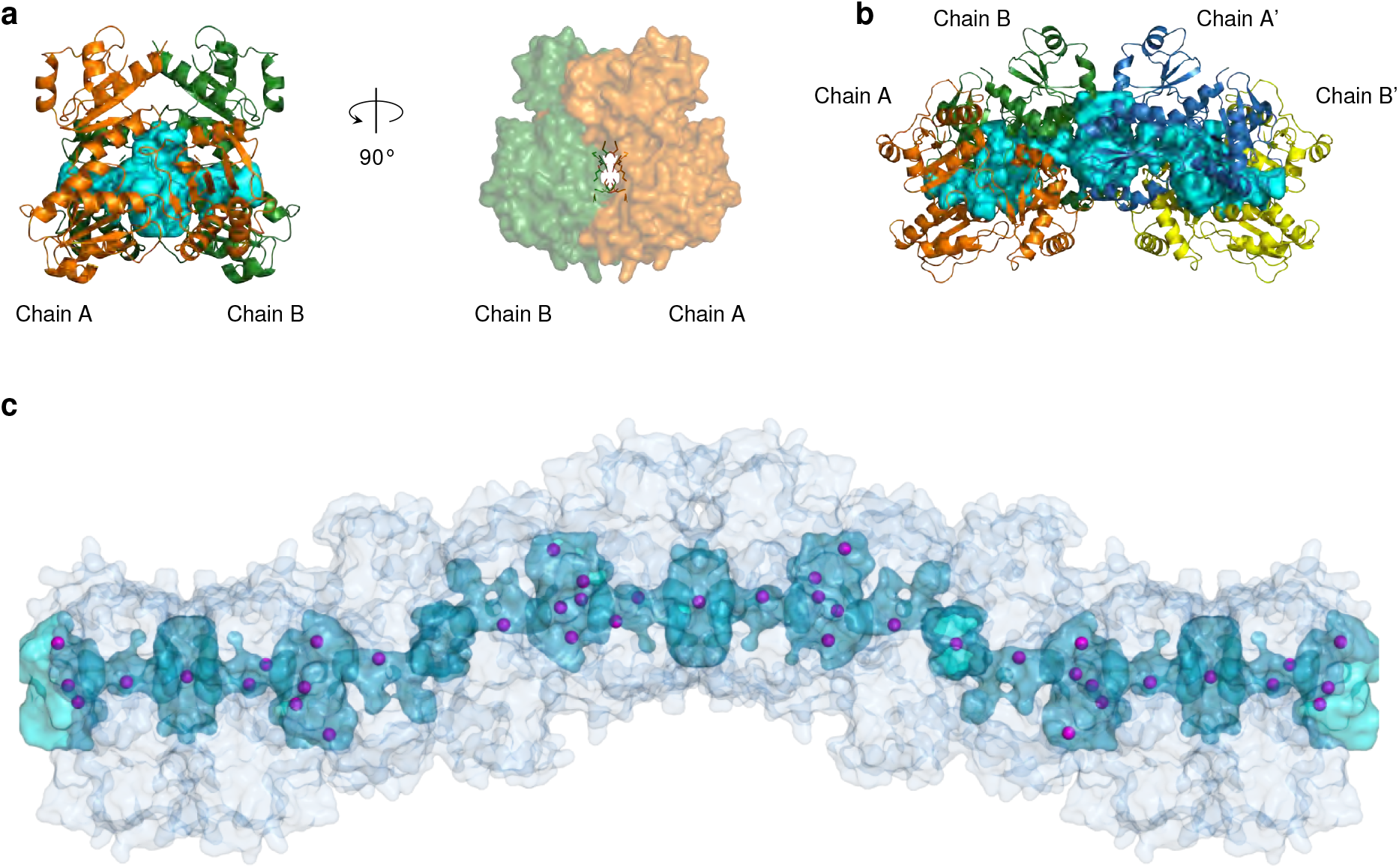
The Cardiac Calsequestrin Filament Possesses a Continuous, Solvent-Accessible Interior Cavity That Winds Along Its Long Axis. **a**, The interior cavity of the dimer (colored cyan) as traced by HOLLOW using a 1.4 Åprobe. Also shown is the interior cavity viewed down its long axis. Residues that participate in the coordination of Yb atoms within the intra-dimer cleft (Figure 4) are shown as sticks. All other residues are rendered as surface. **b**, The lumen of the dimer is continuous with a large solvent pocket formed by the tetramer interface. **c**, View of the filament and its continuous interior cavity, with Yb sites shown as purple spheres.

## Dominant Disease Mutations Disrupt Cardiac Calsequestrin’s Filament-Forming Interface

The newly observed filament structure permits us to assess the possible pathogenic mechanism of the S173I mutation that began this investigation. Remarkably, S173 occupies a critical position at the filament-forming interface - a charged pocket formed by the interaction of K87, S173, and D325 (Figure 7a with supporting electron density map in Figure S9). This pocket, formed of residues from 3 different protomers and also 3 different thioredoxin domains, enforces an interaction between the 3 thioredoxin domains at a single site. This interaction explains the disruptive effect of the S173I mutation discussed above (turbidity assay from Figure 1). On the basis of the apparent importance of this site, as well as the absence of D325 in any other previously described candidate filament interface, we also mutated residue D325 to D325I. As expected, D325I exhibits similarly depressed filamentation kinetics in the turbidity assay (Figure 7b).

**Figure 7:**
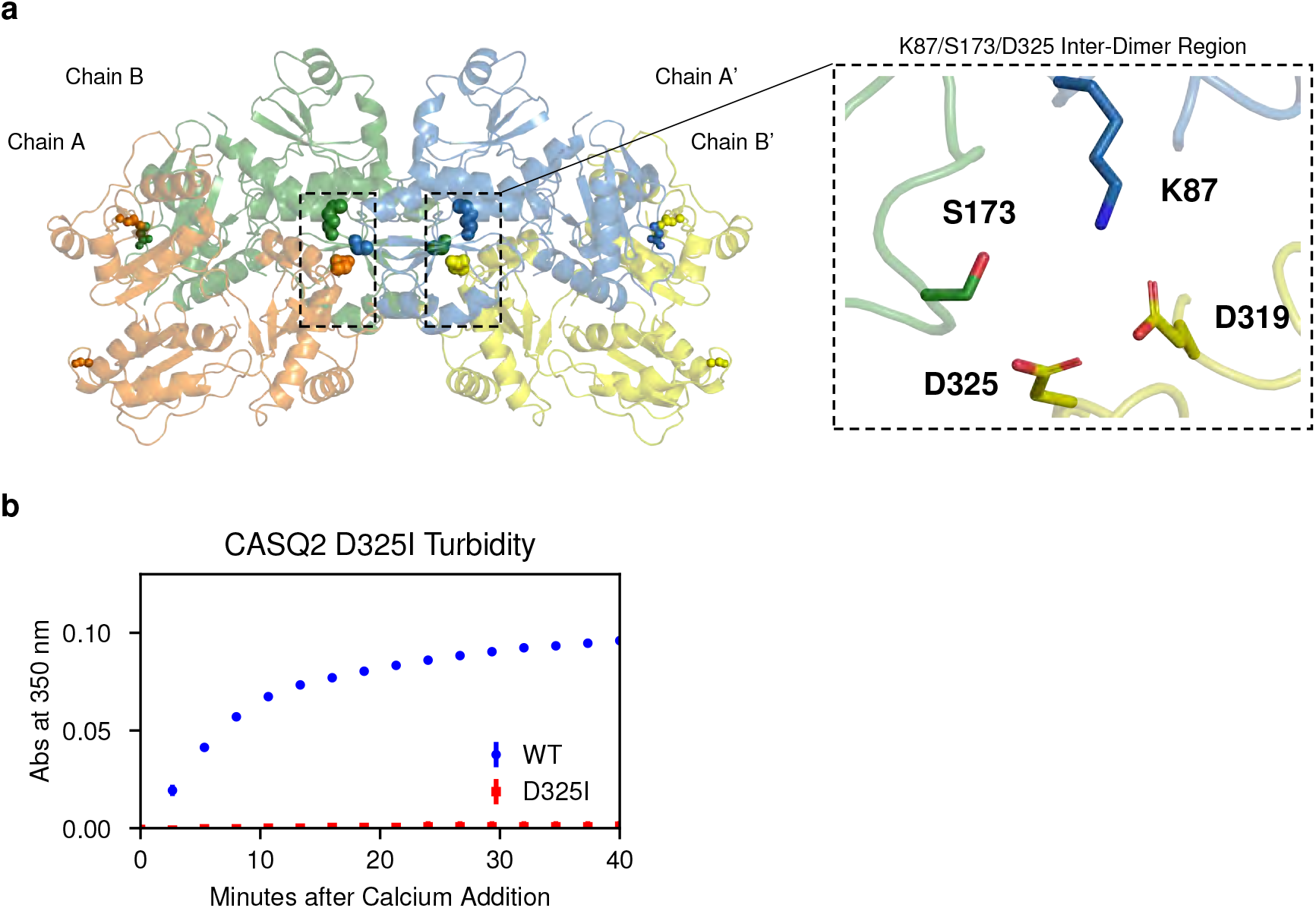
Dominant Disease Mutations Disrupt Cardiac Calsequestrin’s Filament-Forming Interface. **a**, The S173 residue, site of the newly-identified S173I putative disease mutation, is located at a highly hydrophilic 3-protomer interface within the inter-dimer contact region. Disruption of this hydrophilic pocket by hydrophobic substitution leads to filamentation defect. The inter-dimer interface is shown with interface residues in the S173 region rendered as spheres. the closeup panel shows a charged pocket at the location of the CPVT-associated S173I mutation is found. In this pocket, 3 different thioredoxin domains from 3 distinct chains interact (K87, S173, D325). **b**, Turbidity assay with the D325I mutation.

The afore-mentioned K180 residue has recently been implicated in CPVT ***Gray et al. (2016)***. Specifically, a lysine-to-arginine mutation at this position provided the first strong genetic evidence for a dominant CPVT-causing mutation in cardiac calsequestrin. Although this substitution is relatively conservative, the genetic evidence for the pathogenicity of K180R as a dominant mutation is compelling on the basis of a comprehensive segregation analysis in a large family. We have shown in Figure 1 that the K180R mutation also results in a filamentation defect. Our new structure provides a mechanistic explanation for this effect. In the new structure, K180 participates in a helix-stapling salt bridge at the critical E184/E187 inter-dimer Yb site (Figure 5a), shown by us to be essential for robust filamentation. The fact that K180R exhibits filamentation defect only in the presence of magnesium suggests that K180R, rather than disrupting the salt bridge with D50, alters the electrostatics of the nearby cation-binding site. The high conservation of K180 in evolution (Figure S10) and key role of K180 and its interacting partners in supporting calsequestrin filamentation provide additional support for the physiological relevance of our new structure.

## Discussion

We report here a crystal structure of a calsequestrin filament. We also elucidate the biochemical basis of divalent cation-induced filamentation. Using Yb substitution, we confirm previous findings of intra-dimer cation binding sites, and we reveal additional inter-dimer cation sites that have a substantial effect on the rate of multimerization. Furthermore, we provide three forms of evidence in support of the biological relevance of the new structure: biophysical (buried surface area, all-by-all interactions of protomers and domains), biochemical (disruption of multimerization by targeted mutagenesis), and biomedical (disease mutations with dominant inheritance are located at the newly-identified filament-forming interfaces).

What is the proximal source of free energy for calsequestrin filamentation? The prevailing view has been that calsequestrin filamentation is driven by an increase in solvent entropy: ions are bound at filament interfaces, and solvent entropy increases as ion hydration shells are lost ***Krause et al. (1991)***. Under this view, the net contribution from solvent entropy is much greater than the net enthalpic contribution from calcium binding after considering the energetic cost of dehydration. This hypothesis is complicated, however, by the fact that we observe stable filamentation in the near-absence of ligand (only trace concentrations of divalent cations were present in the low-pH crystallization condition). The low-pH filament is stabilized instead by carboxyl-carboxylate interactions between closely juxtaposed acidic residues. While increased solvent entropy likely contributes to filamentation, we postulate instead that protein conformational entropy is likely to be the major driving force. The structure we report is remarkably disordered by conventional B-factor and RSRZ metrics, and especially so in Domain I. In fact, all tightly-packed calsequestrin dimers (7 crystallographic results total, including this study) exhibit increased disorder in the solvent-exposed loops of Domain I (Figure S5). The consistency of this observation within the group makes it less likely that conformational disorder within Domain I is a feature of our specific crystallization condition and more likely that it is a result of the dimer having adopted the tightly-packed conformation favorable for filamentation.

Although low pH may not be necessary for filamentation, calsequestrin’s pH-sensitivity prompts questions about the possible role of intra-luminal pH changes in the regulation of calcium uptake and release. Carboxyl-carboxylate side chain interactions modulated by prevailing pH are a common feature of calcium-binding proteins, wherein a shared proton provides a stabilizing interaction when the cation is unavailable ***Milos et al. (1986);*** Krause et al. (1991). These carboxyl-carboxylate interactions have a higher pKa than a solitary carboxylate, permitting them to play a significant role at pH ranges closer to physiologic ***Sawyer and James (1982);*** Krause et al. (1991). Even at neutral pH, studies of calsequestrin and calmodulin have revealed that protons are released upon calcium binding, consistent with loss of carboxyl-carboxylate bonds ***Milos et al. (1986)***; ***Krause et al. (1991)***. Notably, the SERCA pump, in addition to being an ATPase, is a Ca^2^+/H+ antiporter, so it is reasonable to hypothesize that proton flows over the course of a cycle of CICR have downstream regulatory effects on calsequestrin’s filamentation state, or its affinity for calcium, or both. A small decrease in SR luminal pH during calcium release has been observed ***Kamp et al. (1998)***, and multiple groups have proposed that proton influx into the SR constitutes a small but important fraction of the counterion flow required to maintain charge neutrality when large calcium fluxes occur. Proton efflux from the SR during calcium reuptake, by way of the SERCA pump’s antiporter function, would increase the availability of calcium binding sites. Conversely, protein influx into the SR during calcium release -possibly via a second, independent proton transport pathway within the SERCA pump ***Espinoza-Fonseca (2017)*** – could conceivably stabilize calsequestrin filaments until high levels of calcium are restored. Prior work showed a change in calsequestrin’s intrinsic fluorescence at a pH of 6.0 ***Hidalgo et al. (1996),*** suggesting that dynamic effects on calsequestrin are not limited to the low pH regime used in our crystallization experiments. Since calsequestrin is present at high concentration and likely acts non-trivially as a proton buffer in its own right, effects due to total free proton concentration may manifest as protons move from an SR-based buffer system to one in the cytosol and back with little change in detectable pH. It is important to note that presumed regulatory of effects of dynamic SR pH are speculative and require further elucidation.

Until recently, *CASQ2* disease mutations were thought to be recessively inherited more or less as a rule, with no strong genetic evidence for a dominant disease allele until the report of the K180R mutation ***Gray et al. (2016)***. The recessive inheritance group corresponds to mutations at or near the intra-dimer interface (e.g. R33Q, D307H, and P308L). Since filamentation is necessary for calsequestrin to remain in the jSR ***Milstein et al. (2009)***; ***McFarland et al. (2010)***; ***Knollmann (2010),*** these intradimer interface/filamentation-defective mutants are assumed to be trafficked out of the jSR, leaving any remaining pool of wild type protein largely unaffected. Under this model, filamentation defects overlap mechanistically with a class of calsequestrin-deficient conditions arising from null or hypomorphic alleles, all leading to a final common endpoint of decreased calcium-buffering capacity, and a resulting susceptibility to diastolic calcium leak.

In contrast to the traditional genetic model of CASQ2-associated disease, the mutations investigated in this study (S173I, K180R) are associated with dominant inheritance. These mutations disrupt filamentation at the inter-dimer interface instead of the intra-dimer interface. The apparent discrepancy in inheritance patterns, mapping cleanly to distinct interfaces albeit with limited sampling, is striking. Poisoning of multimerization in classical dominant fashion would seem to require mutant-incorporating dimers to remain jSR-resident, where they could interfere with assembly. There is no reason, however, based on current understanding of secretory pathways, that the dimer would be any less susceptible than the monomer to trafficking out of the jSR. How, then, can the ER export mechanism be reconciled with dominant-acting filamentation-defect mutants? If filamentation defects are equivalent at the molecular level to calsequestrin deficiency, how do we explain the finding of filamentation-disrupting mutations with likely dominant inheritance? We would propose that all filamentation-defective mutants within unincorporated dimers are trafficked out of the jSR in much the same way that unincorporated mutant monomers are. The resulting disease mechanism for dominant mutations is therefore more complex than a classical dominant negative effect: for as long as mutant-containing dimers may be present in the jSR, they interfere with multimerization, but crucially, when they leave, they steal half of the total WT protein with them. The result is paradoxical and perhaps somewhat novel: insufficiency by way of underlying dominant negative biochemistry.

In sum, taking into account calsequestrin’s demonstrated susceptibility to trafficking, we can combine the present work with the existing rich body of calsequestrin research to neatly explain the inheritance puzzle associated with CPVT-causing calsequestrin mutations. Dimer-defective mutants produce monomers which are trafficked away, but the heterozygous state is rescued because WT protein is unaffected by the pool of defective monomeric protein. In contrast, mutations that interfere with the inter-dimer interaction result in depletion of a substantial fraction of WT protein, as unincorporated dimers containing a mix of WT and mutant protein are continually lost. Future work should seek to confirm this hypothetical mechanism using cell biological investigative methods.

## Materials and Methods

### Human Subjects

The patient included in the study provided informed consent as part of a research protocol approved by the University of California, San Francisco Committee on Human Research.

### Cloning and Generation of Plasmids

Full-length cardiac calsequestrin was cloned from human cardiac mRNA by reverse transcription, PCR, A-tailing of a PCR product, and TA ligation. A clone lacking the signal peptide sequence was subcloned by PCR and Gibson Assembly into a T7-based bacterial overexpression vector (pET28a) in front of a 6His site and TEV protease cleavage sequence. Point mutants were generated using the protocol from the Q5 Site-Directed Mutagenesis Kit (New England BioLabs), using either the Q5 or Phusion polymerases. All constructs were transformed in NEB Stable or XL-1 Blue ***E. coli***, purified by miniprep, verified by Sanger sequencing, and retransformed into Rosetta (DE3)pLysS ***E. coli*** for overexpression. Primers used for cloning and mutagenesis are provided in (Table S3). Selection for the pET28a vector was performed using 50 μg/L kanamycin. Selection for pLysS was performed by adding 25 μg/L chloramphenicol.

### Expression and Purification of Cardiac Calsequestrin

pET28a-based expression constructs transformed into Rosetta (DE3)pLysS ***E. coli.*** Overnight starter cultures were used to inoculate large cultures (typically 750 mL of broth per 2.8 L flask), which were grown to OD 0.4 and then induced with 0.25 mM IPTG. Upon induction, temperature was reduced from 37 °C to 24 °C. Cultures were grown for 6-9 hours post-induction or overnight and then spun down (optimal yields were observed from shorter durations of growth). All cultures were grown in standard LB in 50 μg/L kanamycin and 25 μg/L chloramphenicol. Pellets were resuspended in lysis buffer (20 mM Tris pH 7.4, 500 mM NaCl, 10 mM imidazole, 1 EDTA-free protease inhibitor tablet per 50 mL) and frozen at −80 °C. Frozen suspensions were thawed, sonicated on ice (5 min at 1 s on/1 s off), and clarified (15,000 g, 45 min, 4 °C). The clarified supernatant was filtered (0.2 μm), and calsequestrin-containing fractions were isolated by IMAC using a 5 mL HisTrap FF column attached to a GE Akta FPLC (IMAC Buffer A: 20 mM Tris pH 7.4, 500 mM NaCl, 10 mM imidazole; IMAC Buffer B: 20 mM Tris pH 7.4, 500 mM NaCl, 300 mM imidazole). Protein was eluted in 10% steps of Buffer B. The first eluted fraction (10% Buffer B) was always discarded (consistently observed to be impure as determined by SDS-PAGE). Remaining protein-containing fractions were pooled. TEV protease was added to the pooled fractions at a concentration of 1:40 by mass, and the protein was dialyzed overnight at 4 °C in TEV protease dialysis buffer (50 mM Tris pH 8.0, 0.5 mM EDTA, 1 mM DTT). The cleaved protein was further dialyzed for several hours in EDTA dialysis buffer (20 mM HEPES pH 7.3, 100 mM NaCl, 5 mM EDTA) and then overnight into Anion Exchange Buffer A (20 mM HEPES pH 7.3, 100 mM NaCl). Anion exchange polishing was performed using a HisTrap FF column in series with 3×1 mL Mono Q columns. (Buffer A: 20 mM HEPES pH 7.3, 100 mM NaCl; Buffer B: 20 mM HEPES pH 7.3, 1 M NaCl). Protein was eluted in a continuous gradient up to 100% Buffer B), with calsequestrin-rich fractions consistently eluting at 40-50% Buffer B. Fractions were collected and analyzed for purity by SDS-PAGE and A260/A280 ratio. Fractions that were optimally pure and free of A260 contamination were pooled, concentrated to 20 mg/mL, and frozen at −80 °C.

Alanine mutants and the D325I mutant were purified as described above, except that phosphate IMAC buffers were employed (Buffer A: phosphate buffer at pH 7.4, 500 mM NaCl, 10 mM imidazole; Buffer B: phosphate buffer at pH 7.4, 500 mM NaCl, 300 mM imidazole), and an on-column high-salt wash was performed (phosphate buffer at pH 7.4, 2 M NaCl). In addition, the TEV protease dialysis buffer used for these purifications contained 100 mM NaCl.

### Crystallization of Cardiac Calsequestrin

Crystallization screens were carried out in 96-well hanging-drop format and monitored using a Formulatrix Rock Imager automated imaging system. Conditions conducive to crystal growth were optimized and then reproduced in a 24-well format. The best diffraction was obtained by mixing thawed protein (10-20 mg mL^-1^ in 20mM HEPES pH 7.3, 400-500mM NaCl) 1:1 with 15% PEG 4000 and 400mM Li_2_SO_4_. The pH of the PEG 4000 solution used to produce the best-diffracting crystals was tested by litmus paper and found to be approximately 3-3.5. Despite the presence of 20 mM HEPES in the protein reagent, the pH of the drops in which crystals grew was controlled by the PEG and remained 3-3.5. Freshly-made PEGs were incompatible with calsequestrin crystal growth except when concentrated HCl was added to the mother liquor, producing crystals similar to those observed with benchtop-aged PEGs. Interestingly, only unbuffered conditions yielded crystals. Multiple attempts to grow crystals at a buffered low pH (using acetate or glycine-based buffers) failed.

### Ytterbium Soak of Cardiac Calsequestrin Crystals

We initially attempted to identify calcium sites using anomalous signal from calcium (CaCl_2_) added to the crystallization condition described above. We were unsuccessful, likely due to a combination of several factors. The calcium absorption edge is unreachable with conventional tunable x-ray sources and in normal atmosphere; thus, it can only be approached, with resulting weakened anomalous signal. In addition, calsequestrin has an average ***K***_d_ for calcium of 1 mM. Thus, occupancy at a typical site would be expected to be lower as compared to other calcium-binding proteins. Presence of sulfate in the crystallization condition limited calcium concentrations to approximately 14 mM and below, above which a precipitate was observed. Although this limit is above the ***K***_d_, it was insufficient for robust anomalous signal. Crystals of calsequestrin that formed in trace calcium were therefore soaked in YbCl_3_. Hanging drops containing calsequestrin crystals were uncovered and an Eppendorf Microloader was used to inject 2 μL drops (1 μL protein and 1 μL mother liquor) with 200 μL of 2 M YbCl_3_. Data were collected within 5 minutes with no back-soaking.

### Crystal Data Collection and Structure Determination

Hanging drops were uncovered and submerged in a drop of Parabar 10312 (previously known as Paratone). For Yb-soaks, Yb was quickly injected prior to application of the oil. Crystals were looped and pulled through the oil. Excess oil was blotted away, and the loop was mounted directly into the cryostream of the Tom-Alber-Tron endstation at ALS beamline 8.3.1. Frames were collected at 1.116 keV (1.386keV for Yb-soaked crystals) using the endstation’s Pilatus3 S 6M detector using a strategy designed to balance redundancy against radiation damage.

### Structure Determination

Diffraction images were processed with xia2 using the DIALS integration pipeline and a resolution cutoff of CC_1/2_ > 0.3 ***Winter et al. (2018)***; ***Winter (2010)***; ***Evans and Murshudov (2013)***; ***Evans (2006)***; ***Winn et al. (2011)*** For the native structure, the merged diffraction intensities were used to find a molecular replacement solution in Phaser ***McCoy et al. (2007)*** with the previously published canine cardiac calsequestrin structure, 1SJI ***Park et al. (2004),*** serving as the initial model. This resulted in a solution in space group P4_3_22 containing one calsequestrin chain per AU. This solution was refined in PHENIX ***Adams et al. (2010)*** with PHENIX AutoBuild ***Adams et al. (2010)***; ***Afonine et al. (2012)***; Terwilliger (2004); ***Terwilliger et al. (2008)***; ***Zwart et al. (2005),*** with extensive manual model-building in Coot ***Emsley et al. (2010)***. For the Yb-complexed dataset, data were processed as above but with preservation of anomalous signal (no merging of Friedel pairs). The refined native structure was used a molecular replacement model, and a solution was found in space group P4_3_2_1_2 containing a dimer in the AU. This solution refined poorly. The Yb-complexed dataset was reprocessed in P1, and a molecular replacement solution was found in P1 with 16 chains in the AU. Refinement of this model was tested using Zanuda ***Lebedev and Isupov (2014),*** and the best R-free was found to be in space group P12_1_1. The Yb-complexed dataset was reprocessed in space group P12_1_1, and a molecular replacement solution was found with 8 chains in the AU. This solution refined well. The anomalous map was used in refinement to place Yb atoms in the structure.

### Turbidity Assays

Recombinant protein samples were thawed, diluted in 2 mL-3 mL of Turbidity Assay Buffer (15 mM Tris pH 7.4, 20 mM NaCl, 85 mM KCl) and dialyzed in Turbidity Assay Buffer plus 10 mM EDTA. Samples were then re-dialyzed overnight in the same buffer without EDTA. Approximately half of the sample was then dialyzed in Zero-Potassium Turbidity Assay Buffer (15 mM Tris pH 7.4, 20 mM NaCl) to allow for turbidity results in a zero-potassium context. Protein A280 was measured in triplicate (Nanodrop) using the appropriate matching buffer as background, and protein was diluted to 2.25 μM in a 140 μL volume in half-area wells of a μClear 96-well plate. The plate was covered, and protein in the wells was allowed to equilibrate on the benchtop for 20 minutes. The turbidity assay was performed using a BioTek Synergy 2 plate reader equipped with reagent injectors. Seven μL of 20 mM CaCl_2_ solution was injected into each well, and the plate underwent shaking for 20 s. Absorbance at 350 nm was monitored for 45 min. The protocol was performed in plate synchronized mode for consistent well-to-well timing. A 100 mM ion-selective electrode calcium standard (Sigma, cat no. 21059) stock solution was used for all CaCl_2_ dilutions.

### Quantification and Statistical Methods

Data points in figures represent mean values, with error bars representing standard deviation. All turbidity assay data points are mean of 3 technical replicates.

### Data and Software Availability

The structures determined as part of this work are deposited in the Protein Data Bank (PDB) under identifiers 6OWV (native) and 6OWW (ytterbium-soaked). The raw diffraction dataset for the native structure is deposited in Zenodo under doi:10.5281/zenodo.2941360. The raw diffraction dataset for the ytterbium-complexed structure is likewise deposited in Zenodo under doi:10.5281/zenodo.2943248. Protein structure figures were generated using PyMOL ***Schrödinger, LLC (2015)***. The interior cavity of the calsequestrin filament was traced using HOLLOW ***Ho and Gruswitz (2008)***. Sequence alignments were generated using TEXshade ***Beitz (2000)***. Plots were generated using python mat-plotlib ***Hunter (2007)***. The manuscript and all figure layouts were constructed entirely in LTEX using PGF/TikZ. Data and code to generate all figures are freely available at https://github.com/errontitus/casq2-structure-function.

## Acknowledgments

We thank Jason Roberts for expert clinical review. We thank Christopher Agnew, Jamie Fraser, James Holton, Lijun Liu, Marco Lolicato, Bruk Mensa, and Tarjani Thaker for technical assistance with crystallographic data collection and processing. We thank Lijun Liu and Tarjani Thaker for technical assistance with protein preparation and crystallization screens. We thank Bruk Mensa for technical assistance with plate reader instrumentation. We thank Michael Grabe, Aimee Kao, Dan Minor, and Oren Rosenberg for helpful discussions.

Beamline 8.3.1 at the Advanced Light Source is operated by the University of California Office of the President, Multicampus Research Programs and Initiatives grant MR-15-328599 the National Institutes of Health (R01 GM124149 and P30 GM124169), Plexxikon Inc., and the Integrated Diffraction Analysis Technologies program of the US Department of Energy Office of Biological and Environmental Research. The Advanced Light Source (Berkeley, CA) is a national user facility operated by Lawrence Berkeley National Laboratory on behalf of the US Department of Energy under contract number DE-AC02-05CH11231, Office of Basic Energy Sciences.

E.W.T. was previously supported by a Sarnoff Foundation Fellowship and is currently supported by NIH/NHLBI F30 grant F30HL137329 and NIH/NIGMS grant T32GM007618 to the UCSF Medical Scientist Training Program (MSTP). This work was supported in part by NIH/NHLBI grant DP2HL123228 (to R.C.D.) as well as American Heart Association grant 17IRG33460152 (to R.C.D.).

## Author Contributions

R.C.D. and E.W.T conceived and designed the study. R.C.D. and E.W.T. designed and oversaw all experiments. E.W.T., F.H.D., and C.S. performed experiments. M.S. and J.W. collected and analyzed clinical data. E.W.T. analyzed experimental data. E.W.T. wrote the manuscript. M.S., J.W., N.J., and R.C.D. reviewed and edited the manuscript.

## Declaration of Interests

The authors declare no competing interests.

---

**Figure S1:**
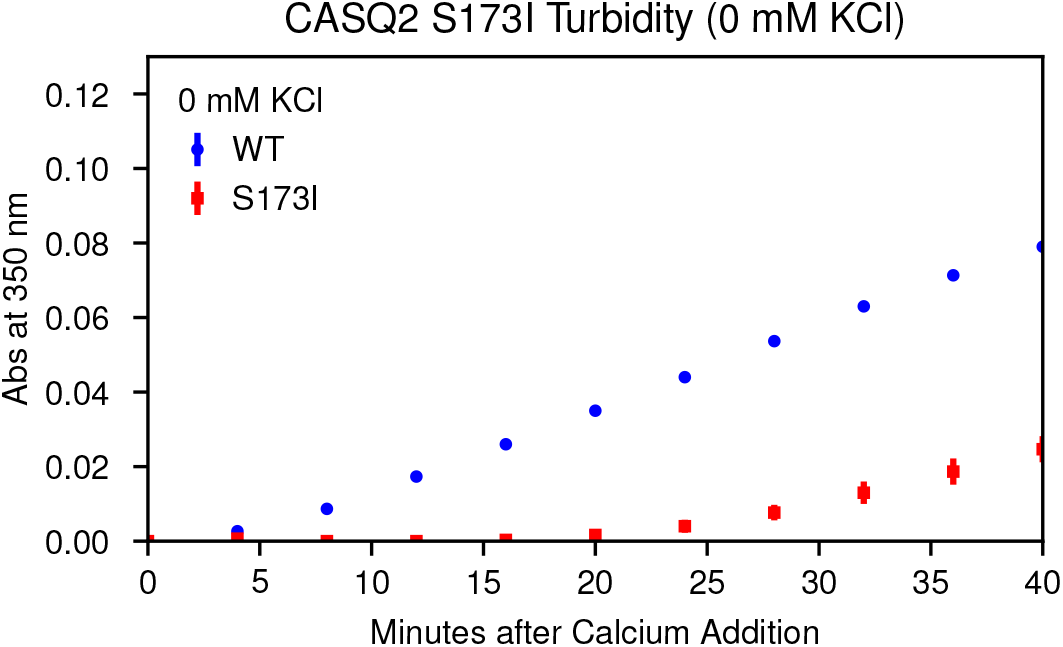
Multimerization Kinetics of the S173I Mutant Observed in 0 mM KCl, Related to Figure 1.

**Figure S2:**
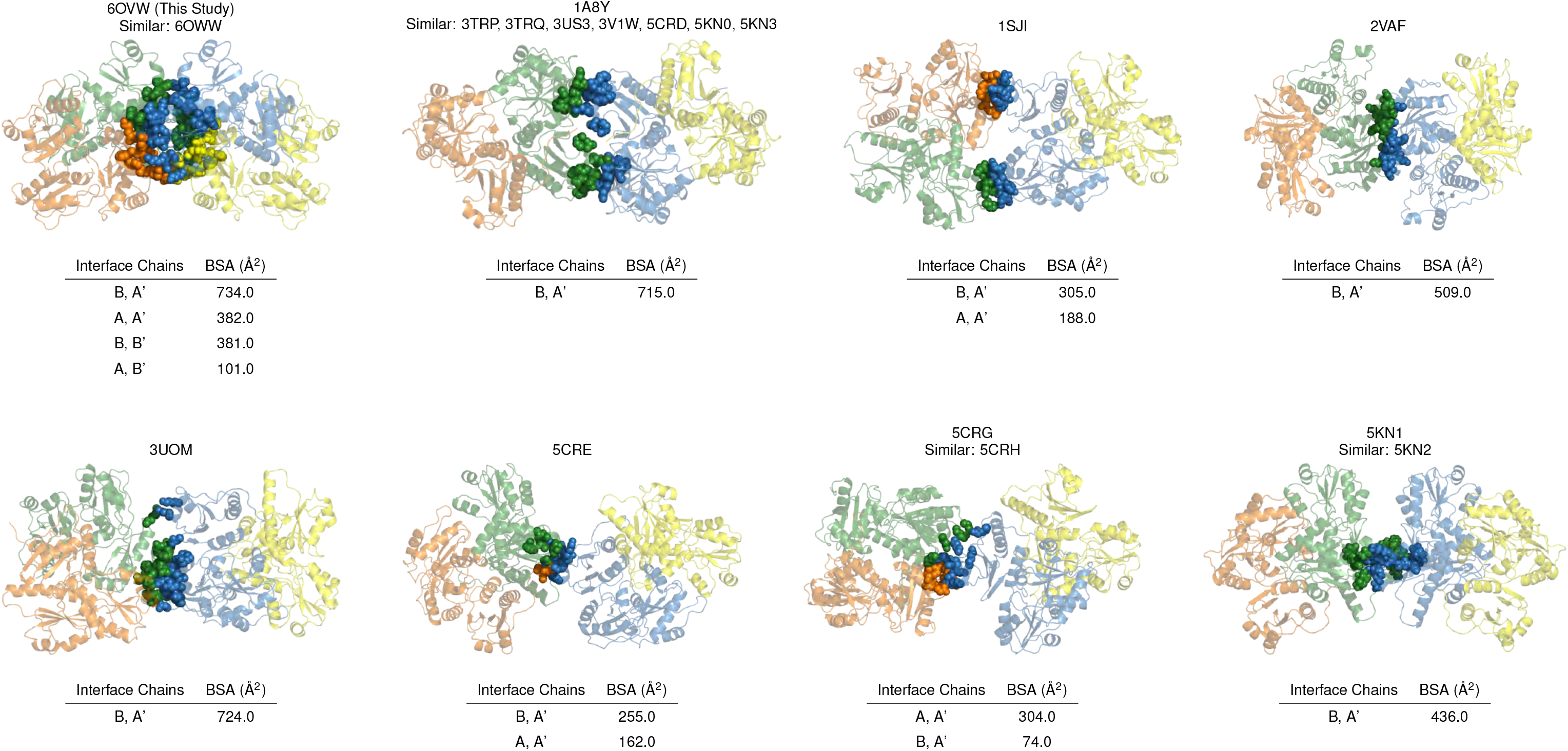
The Dimer-to-Dimer Interface of the New Candidate Cardiac Calsequestrin Filament Is Characterized by All-by-All Contacts and Greater Buried Surface Area (BSA) Compared to All Other Published Calsequestrin Structures, Related to Figure 2. For each published calsequestrin structure, the dimer-to-dimer interface with the greatest buried surface area is shown. Residues with buried surface area at the interface are rendered as spheres. Orange, chain A; forest, chain B; sky blue, chain A’; yellow, chain B’. Where similar PDB codes are listed, dimer-to-dimer interfaces are roughly are isomorphous with the example structure shown, although the space group and unit cell used to determine the structure sometimes differ.

**Figure S3:**
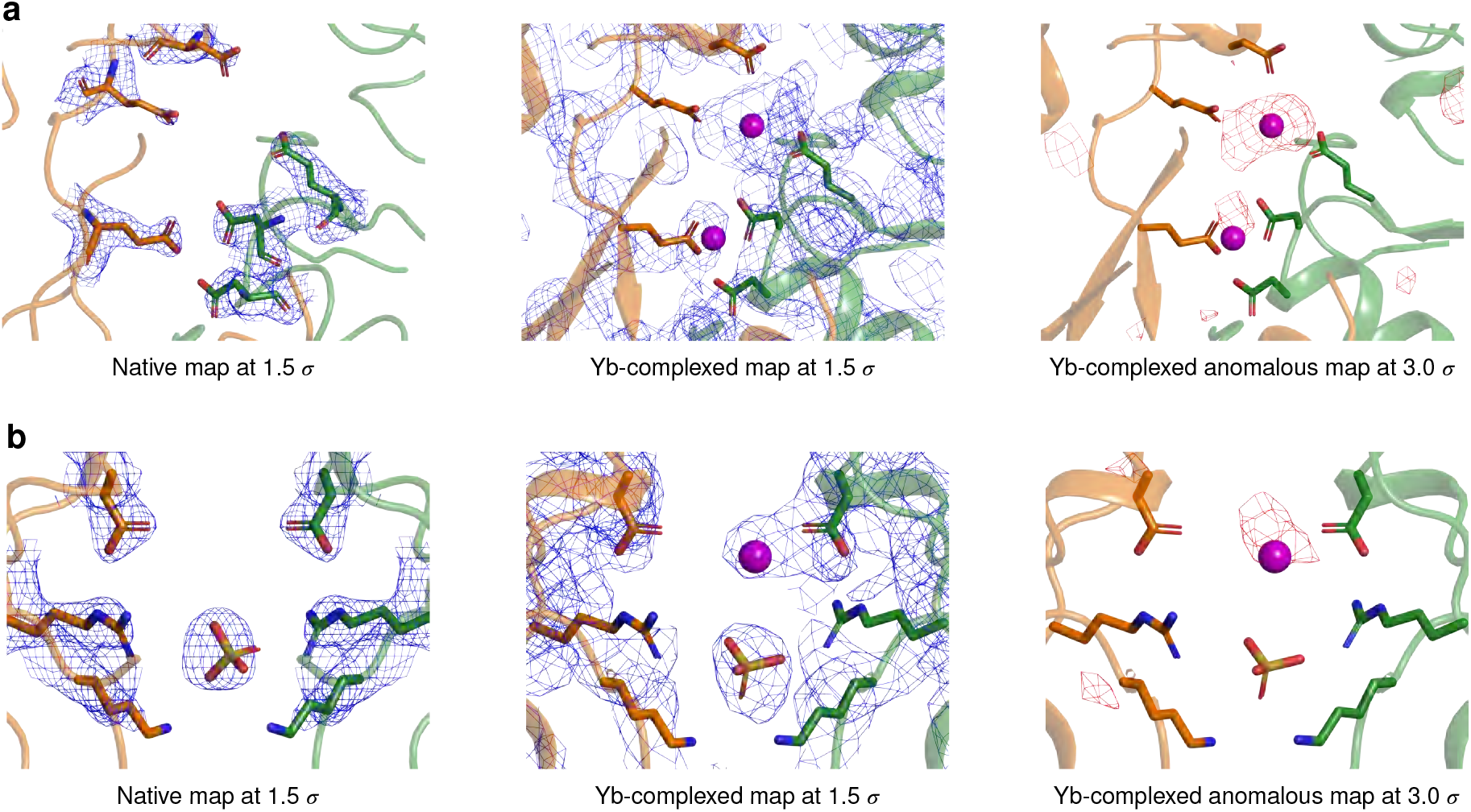
Electron Density and Anomalous Difference Maps for Yb-Binding Sites at the Cardiac Calsequestrin Intra-Dimer Interface, Related to Figure 4. **a**, Electron density and anomalous difference Maps for the D140/E143/E147 region of interest. **b**, Electron density and anomalous difference maps for the D310 region of interest.

**Figure S4:**
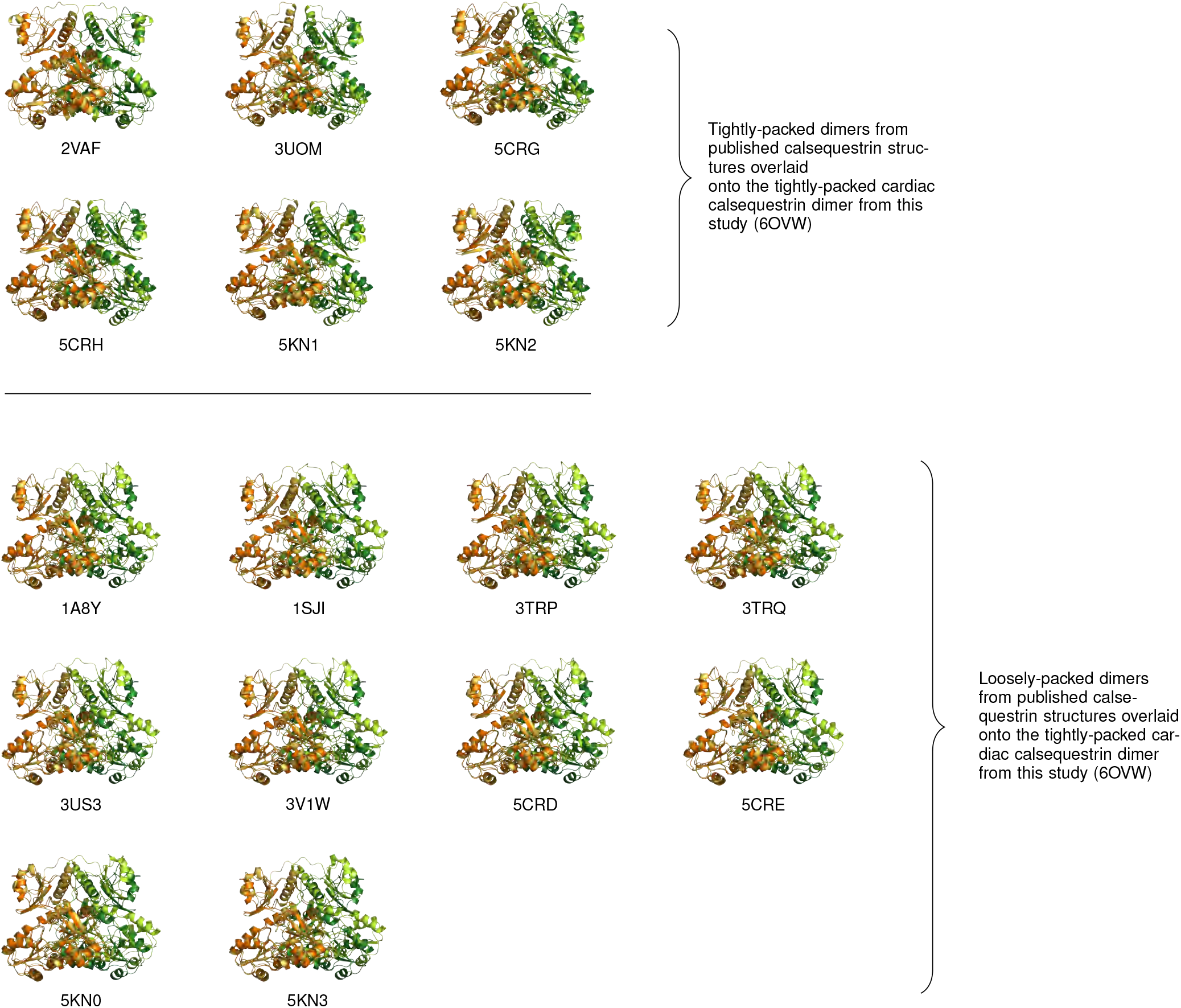
Dimer Overlays Reveal That All Published Calsequestrin Structures Can Be Classified into Tightly-Packed or Loosely-Packed Dimers, Related to Figure 4. Dimers from published calsequestrin structures (lighter orange and green) are overlaid onto the tightly-packed dimer from this study (6OVW, darker orange and green). In each dimer pair, chain A is aligned to chain A to illustrate the relative displacement of chain B. The overlays reveal two distinct conformational groupings. The more tightly-packed conformation with inwardly-rotated chains resembles the dimer in this study. This conformation forms at low pH or in the presence of neutralizing multivalent cations.

**Figure S5:**
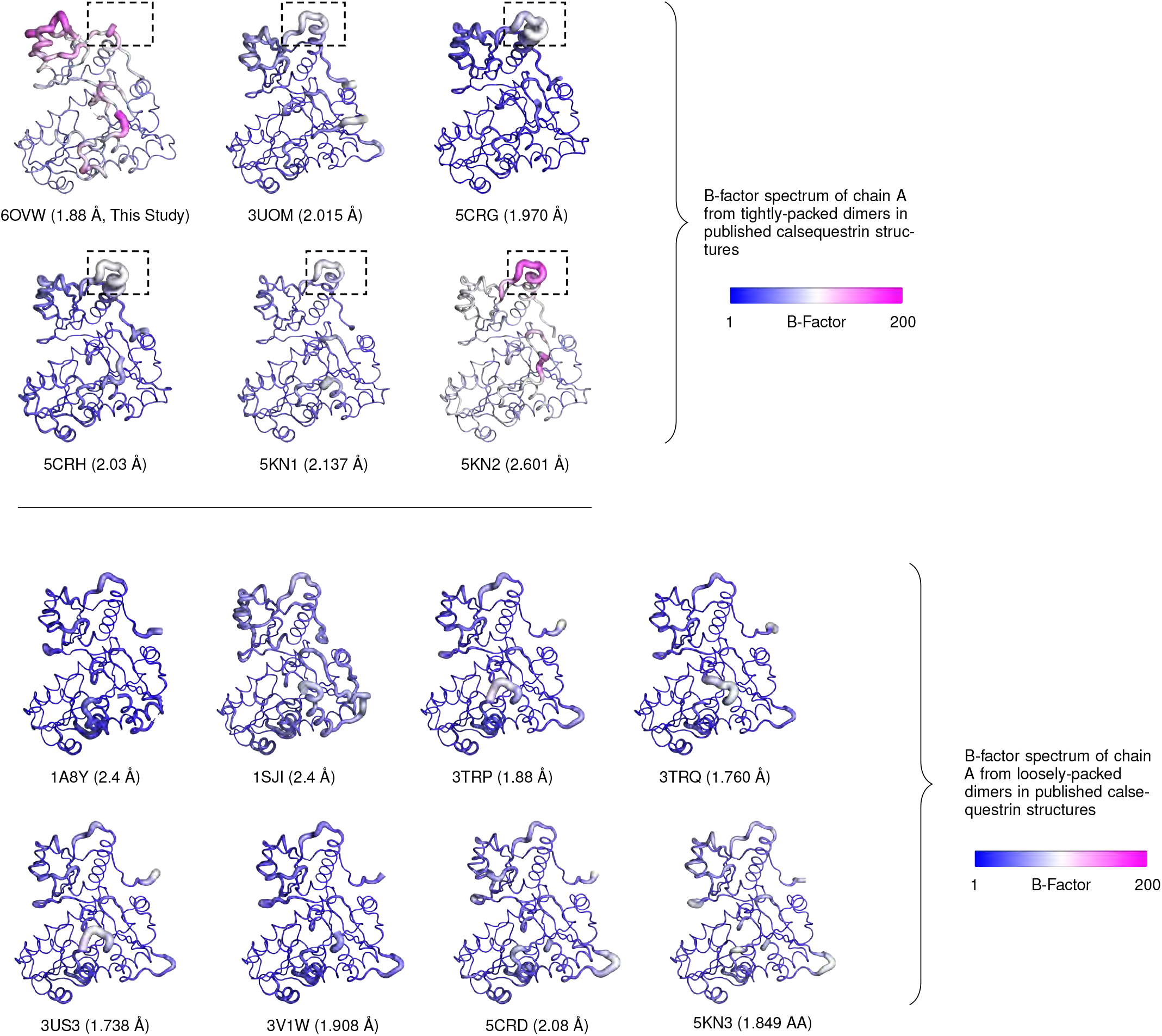
Tightly-Packed Calsequestrin Dimers Consistently Exhibit Increased Conformational Disorder in Domain I, Related to Figure 4. The top panel shows tightly-packed calsequestrin dimers (i.e. dimers of calsequestrin crystallized in low pH or with high concentration of multivalent cations). In these structures, solvent-exposed loops in domain I are consistently disordered. In PDB 6OVW, corresponding to this study, the disordered loop region is omitted entirely due to the high level of disorder. This same region (boxed, residues 58-68) is highly disordered in similar structures. The bottom panel shows loosely-packed calsequestrin dimers (i.e. dimers of calsequestrin crystallized at neutral pH with low or trace concentrations of multivalent cations). The resolution for each structure is indicated, and several structures of non-comparable resolution are excluded (2VAF, 5CRE, 5KN0).

**Figure S6:**
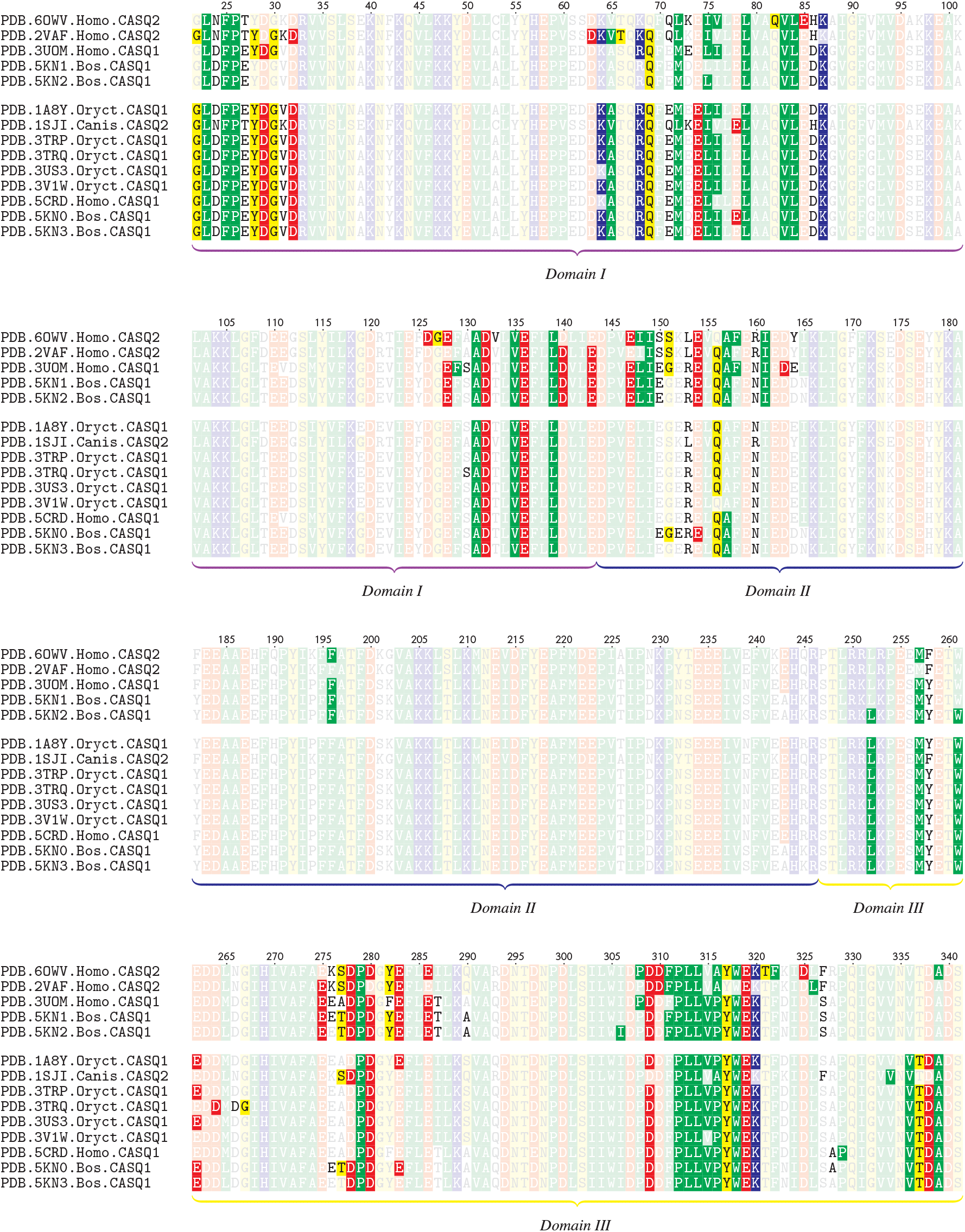
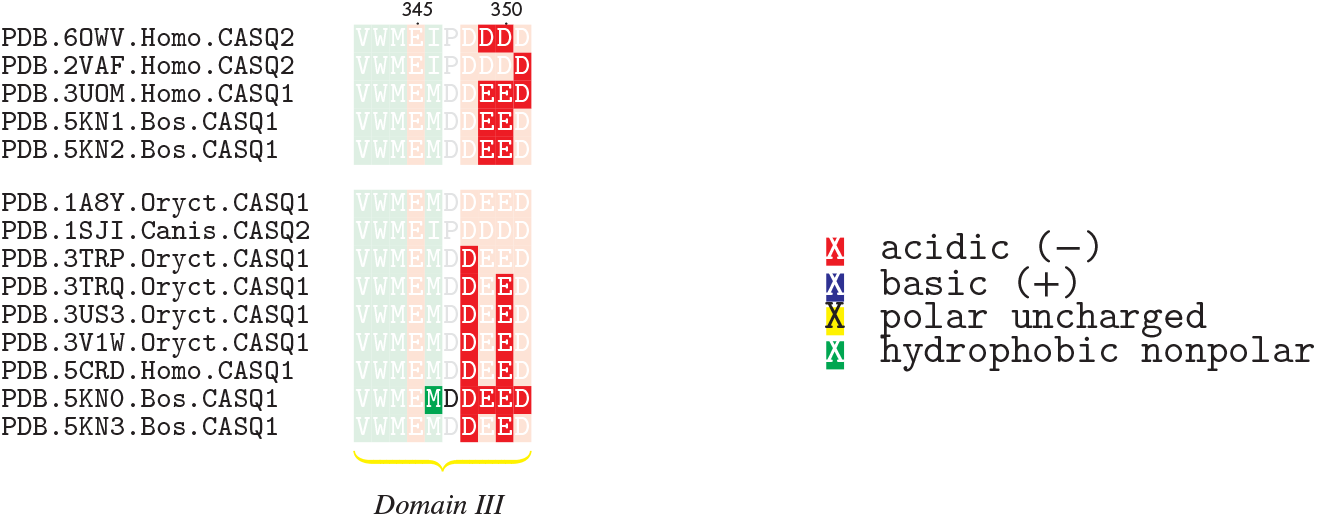
Multiple Sequence Alignment Comparing Intra-Dimer Interface Residues from Published Calsequestrin Structures, Related to Figure 4. Intra-dimer interface residues are highlighted; color represents hydropathy. Alignment is grouped by dimer conformational class (top group: tightly-packed dimers; bottom group: loosely-packed dimers). Rotation of chains in the tightly-packed dimers leads to loss of contacts near the N terminus but gain of contacts elsewhere. Calsequestrin structures 5CRE, 5CRG, and 5CRH are omitted (point mutants belonging to the same investigation as 5CRD).

**Figure S7:**
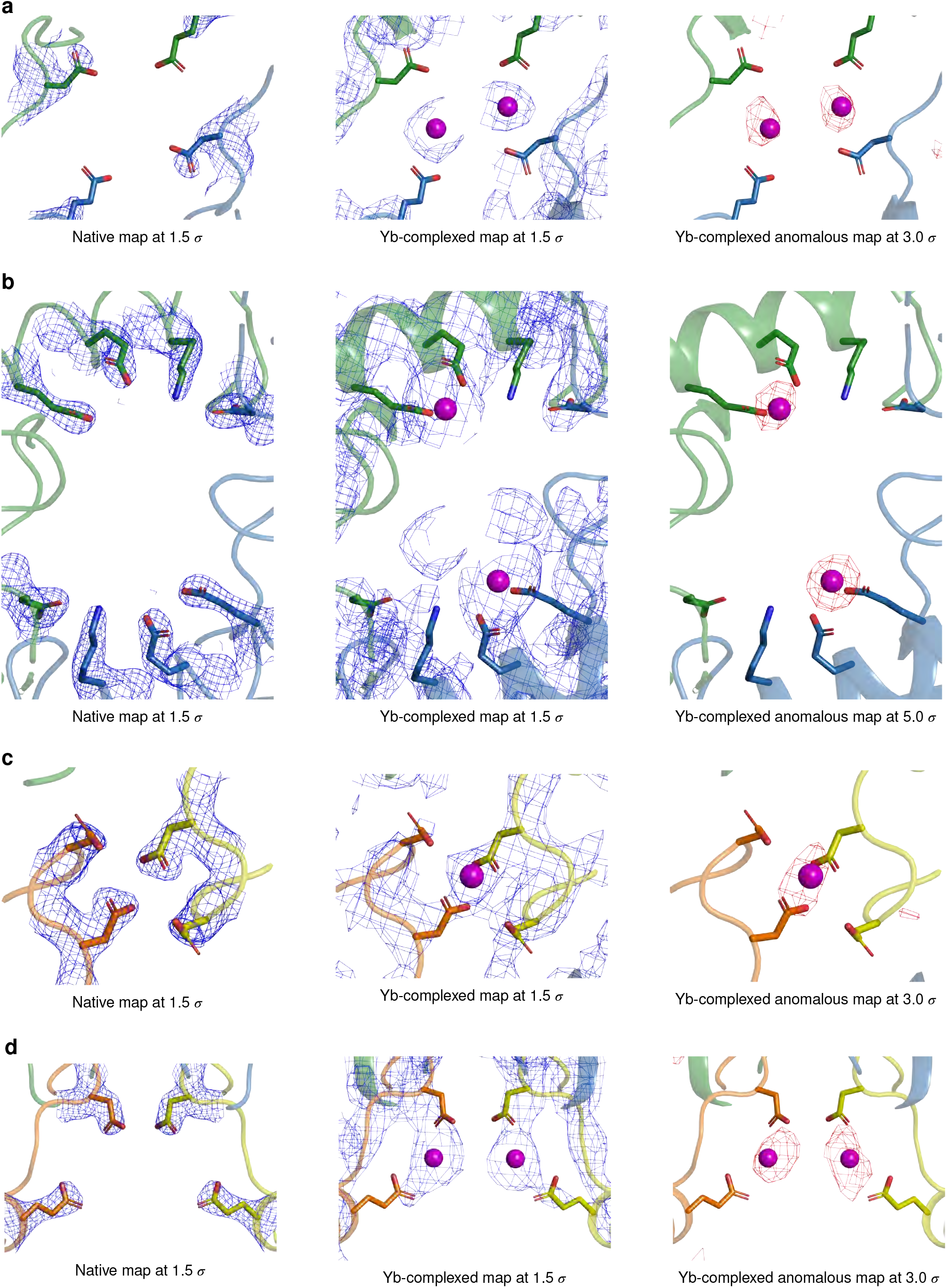
Electron Density and Anomalous Difference Maps for Yb-Binding Sites at the Cardiac Calsequestrin Filament’s Inter-Dimer Interface, Related to Figure 5. **a**, Electron density and anomalous difference maps for the D144/E174 region of interest. **b**, Electron density and anomalous difference maps for the D50/K180/E184/E187 region of interest. **c**, Electron density and anomalous difference maps for the D348/D350 region of interest. **d**, Electron density and anomalous difference maps for the D351/E357 region of interest.

**Figure S8:**
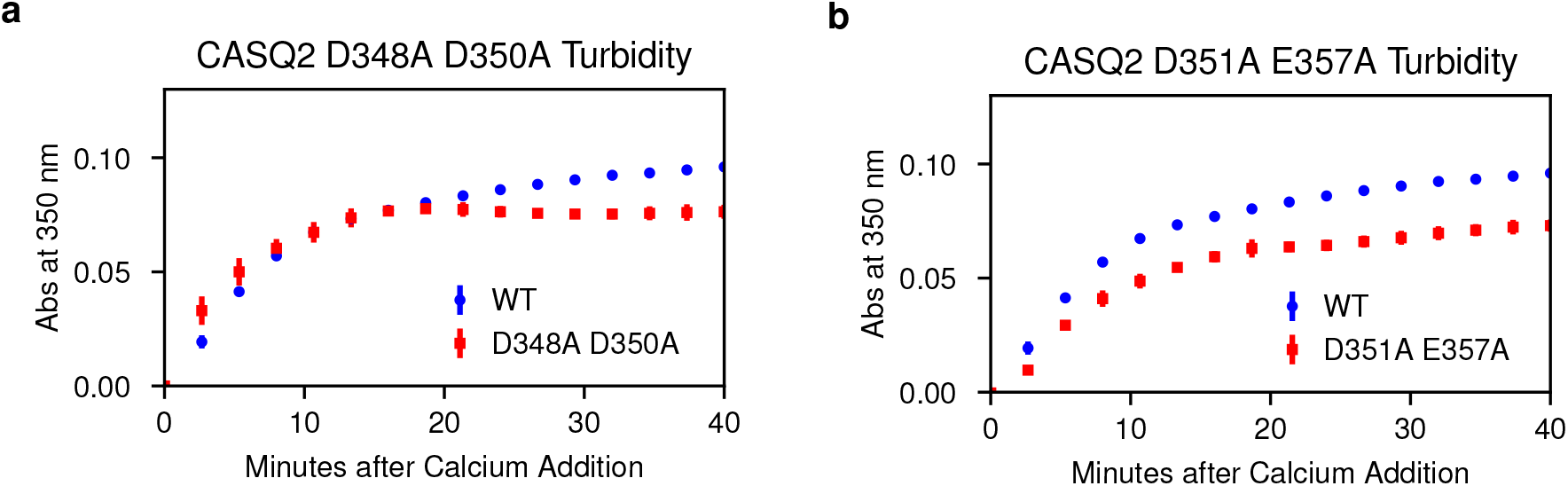
Turbidity Assays Showing Effect of Alanine Mutagenesis of Additional Yb-Binding Sites at the Cardiac Calsequestrin Inter-Dimer Interface, Related to Figure 5. **a**, Turbidity assay after alanine mutagenesis of the putative calcium-coordinating residues D348 and D350. **b**, Turbidity assay after alanine mutagenesis of the putative calcium-coordinating residues D351 and E357.

**Figure S9:**
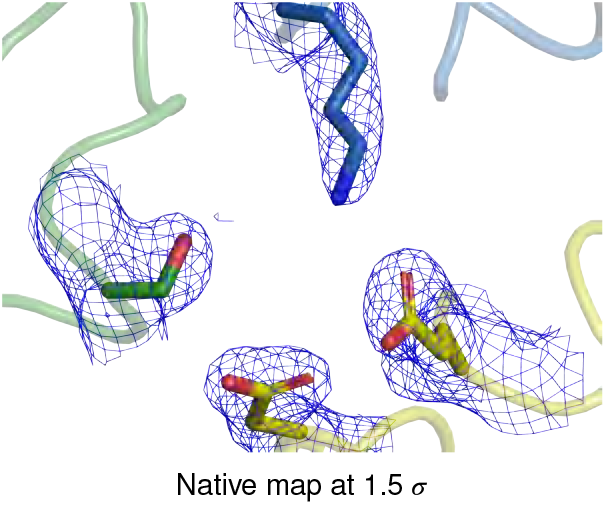
Electron Density Map for the S173 Region at the Cardiac Calsequestrin Filament’s Inter-Dimer Interface, Related to Figure 7.

**Figure S10:**
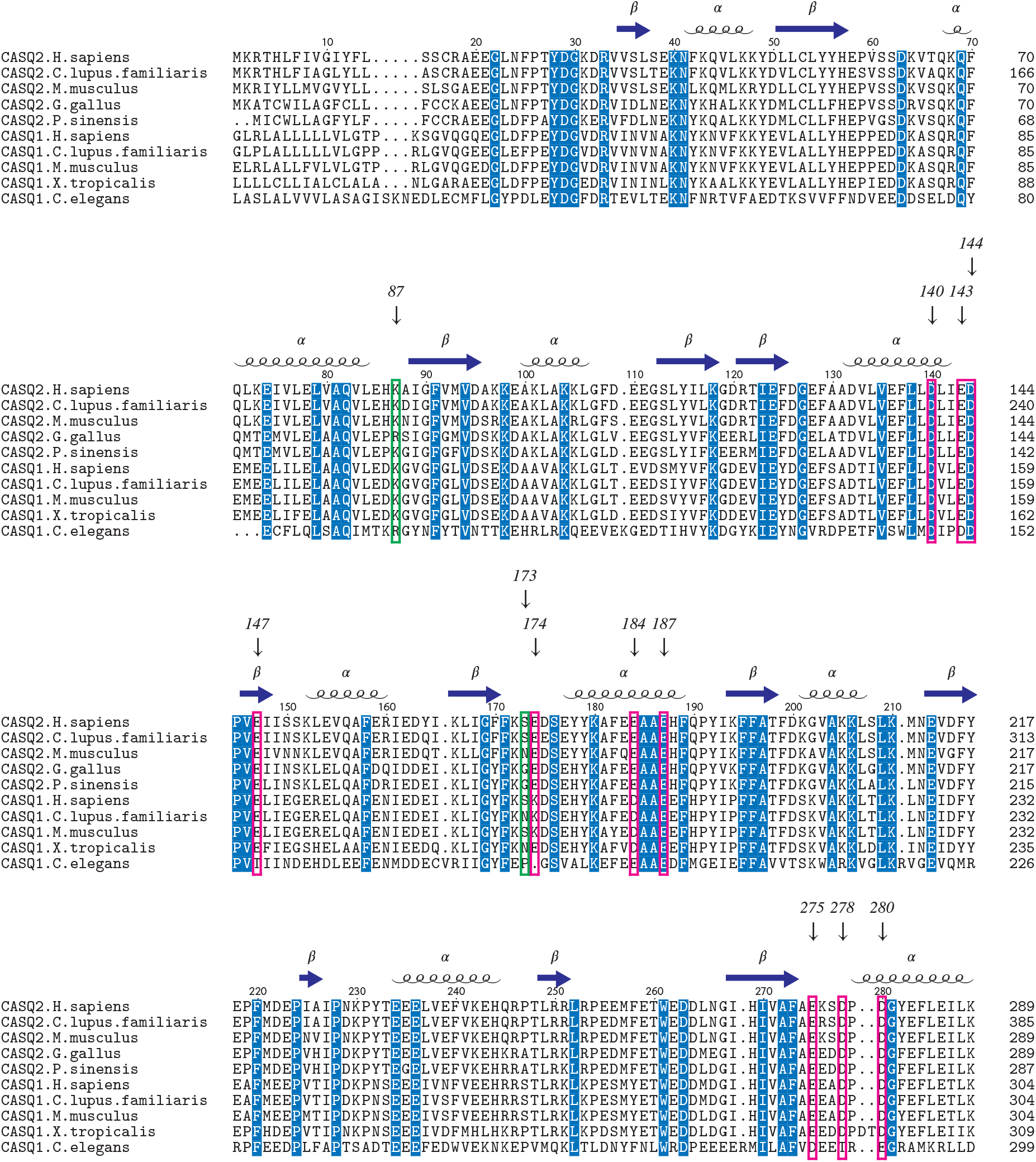
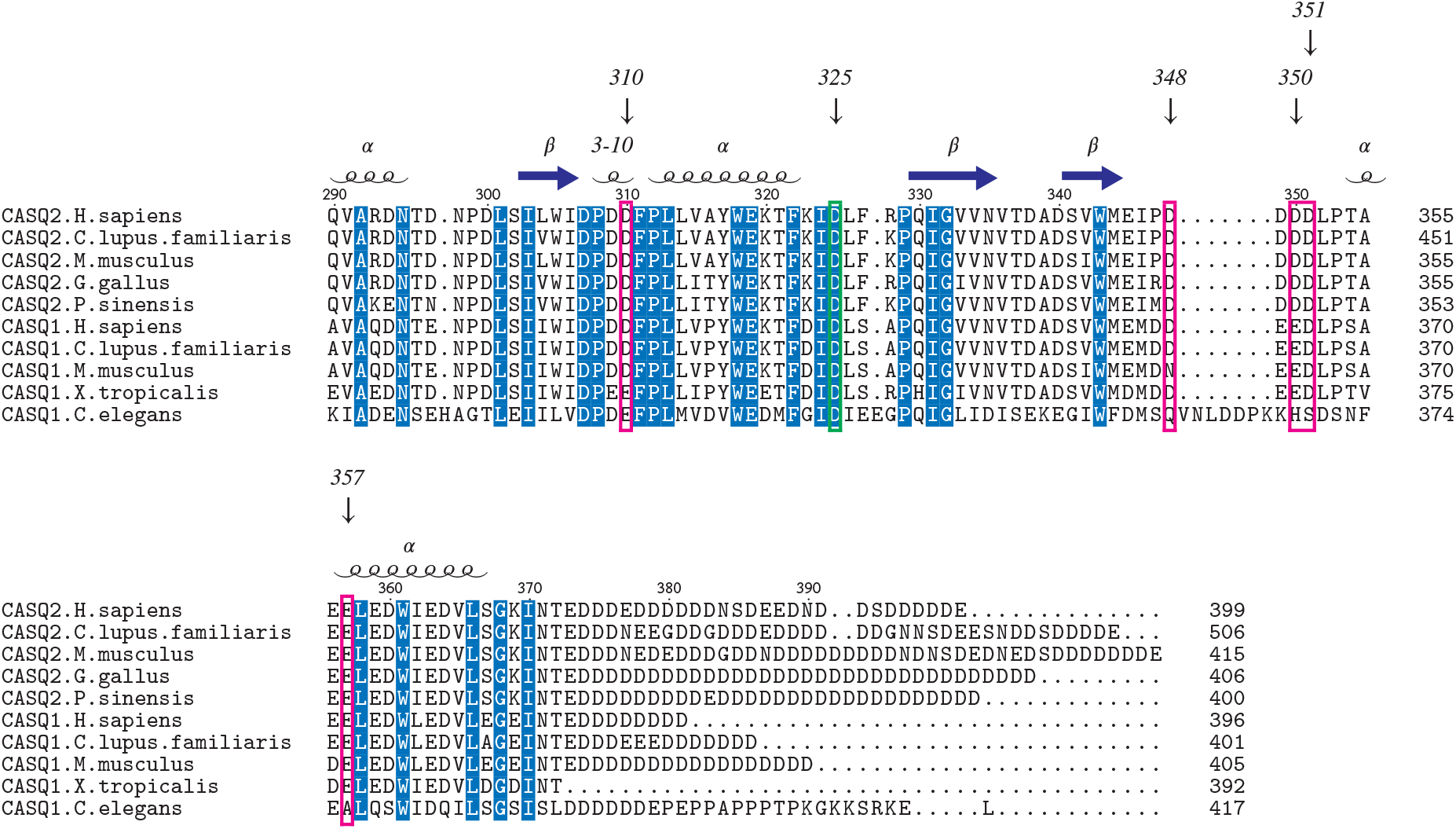
Multiple Sequence Alignment Showing Conservation Across Calsequestrins. Residues of importance at intra-dimer and inter-dimer interfaces are indicated. Magenta boxes: residues implicated in Yb coordination, related to Figure 4 and Figure 5. Green boxes: residues at the 3-protomer dimer-dimer interface, related to Figure 7.

## Supplemental Tables

**Table S1:** Crystallographic Data Collection and Refinement

|  | Native (6OWV) | Ytterbium-Complexed (6OWW) |
| --- | --- | --- |
| **Data Collection** |  |  |
| Wavelength (Å) | 1.116 | 1.386 |
| Space Group | P 43 2 2 | P 1 21 1 |
| Cell Dimensions |  |  |
| a, b, c (Å) | 62.5329, 62.5329, 213.189 | 85.8318, 86.0152, 214.34 |
| $\alpha, \beta, \gamma$ (°) | 90, 90, 90 | 90, 89.9072, 90 |
| Resolution (Å) | 53.94 - 1.88 (1.947 - 1.88) | 107.2 - 3.84 (3.977 - 3.84) |
| $R_{\text{pim}}$ (%) | 1.8 (194.8) | 16.2 (85.0) |
| CC<sub>1/2</sub> | 0.999 (0.422) | 0.992 (0.616) |
| Mn(I)/$\sigma$(I) | 14.82 (0.52) | 4.52 (0.96) |
| Completeness (%) | 97.76 (79.77) | 98.45 (98.05) |
| Multiplicity | 22.5 (12.6) | 11.2 (11.5) |
| No. Unique Reflections | 35501 (3425) | 29897 (2924) |
| **Refinement** |  |  |
| $R_{\text{work}} / R_{\text{free}}$ (%) | 21.4 / 25.4 | 29.1 / 34.2 |
| Resolution (Å) | 53.94 - 1.88 | 107.2 - 3.84 |
| No. of chains in AU | 1 | 8 |
| Number of non-hydrogen atoms | 2779 | 21752 |
| Protein | 2718 | 21665 |
| Ligand | 27 | 87 |
| Solvent | 34 | N/A |
| B-Factors |  |  |
| Protein | 81.93 | 97.36 |
| Ligand | 112.54 | 135.46 |
| Solvent | 55.84 | N/A |
| R.m.s. deviations |  |  |
| Bond lengths (Å) | 0.009 | 0.008 |
| Bond angles (°) | 1.25 | 1.14 |

**Table S2:** Comparison of Crystallographic Space Group and Unit Cell Across All Published Calsequestrin Structures, Related to Figure 2

| PDB Code | Year | Protein | Space Group | Unit Cell Edges | Unit Cell Angles |
| --- | --- | --- | --- | --- | --- |
| 6OWV | 2019 | CASQ2 (Homo) | P 43 2 2 | 62.52, 62.52, 213.15 | 90, 90, 90 |
| 6OWW | 2019 | CASQ2 (Homo) | P 1 21 1 | 85.8318, 86.0152, 214.34 | 90, 89.9072, 90 |
| 1A8Y | 1998 | CASQ1 (Oryctolagus) | C 2 2 21 | 59.740, 145.560, 111.790 | 90.00, 90.00, 90.00 |
| 1SJI | 2004 | CASQ2 (Canis) | I 4 | 145.188, 145.188, 99.82 | 90, 90, 90 |
| 2VAF | 2007 | CASQ2 (Homo) | I 41 2 2 | 150.650, 150.650, 227.470 | 90.00, 90.00, 90.00 |
| 3TRP | 2012 | CASQ1 (Oryctolagus) | C 2 2 21 | 59.122, 144.863, 110.376 | 90.00, 90.00, 90.00 |
| 3TRQ | 2012 | CASQ1 (Oryctolagus) | C 2 2 21 | 59.271, 144.565, 111.170 | 90.00, 90.00, 90.00 |
| 3UOM | 2012 | CASQ1 (Homo) | P 1 | 89.799, 89.793, 119.161 | 90.13, 89.90, 60.05 |
| 3US3 | 2012 | CASQ1 (Oryctolagus) | C 2 2 21 | 59.214, 144.811, 110.471 | 90.00, 90.00, 90.00 |
| 3V1W | 2012 | CASQ1 (Oryctolagus) | C 2 2 21 | 59.082, 144.592, 110.965 | 90.00, 90.00, 90.00 |
| 5CRD | 2015 | CASQ1 (Homo) | C 2 2 21 | 59.170, 145.132, 110.242 | 90.00, 90.00, 90.00 |
| 5CRE | 2015 | CASQ1 (Homo) | P 21 21 2 | 66.106, 82.815, 89.269 | 90.00, 90.00, 90.00 |
| 5CRG | 2015 | CASQ1 (Homo) | P 1 21 1 | 91.179, 67.462, 158.062 | 90.00, 96.48, 90.00 |
| 5CRH | 2015 | CASQ1 (Homo) | P 1 21 1 | 65.681, 68.553, 99.262 | 90.00, 92.84, 90.00 |
| 5KN0 | 2016 | CASQ1 (Bos) | P 1 | 60.342, 92.994, 101.849 | 71.12, 84.57, 73.48 |
| 5KN1 | 2015 | CASQ1 (Bos) | C 2 2 21 | 135.669, 165.604, 156.626 | 90.00, 90.00, 90.00 |
| 5KN2 | 2016 | CASQ1 (Bos) | C 2 2 21 | 130.363, 169.194, 155.477 | 90.00, 90.00, 90.00 |
| 5KN3 | 2016 | CASQ1 (Bos) | C 2 2 21 | 59.393, 146.060, 110.340 | 90.00, 90.00, 90.00 |

**Table S3:**
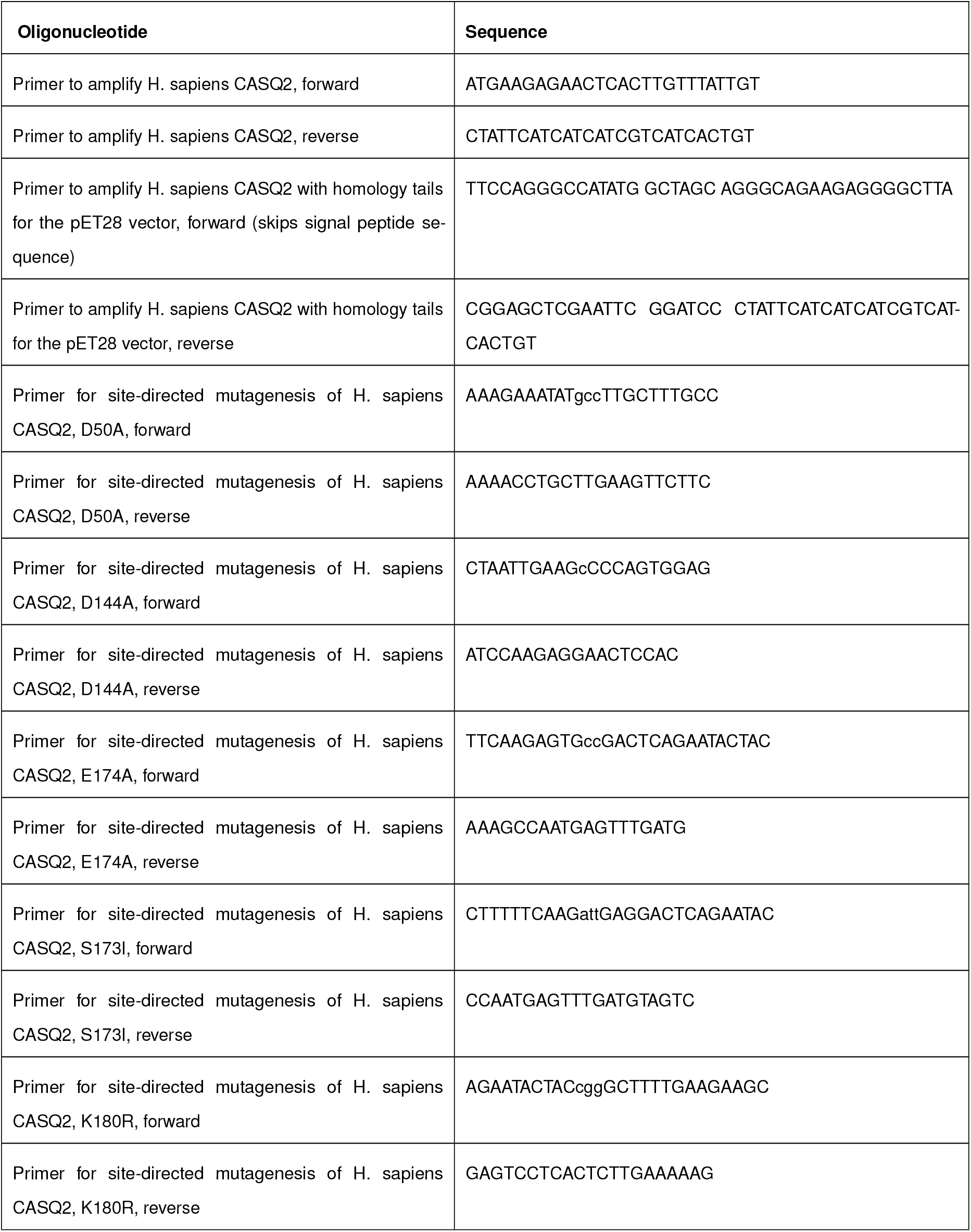
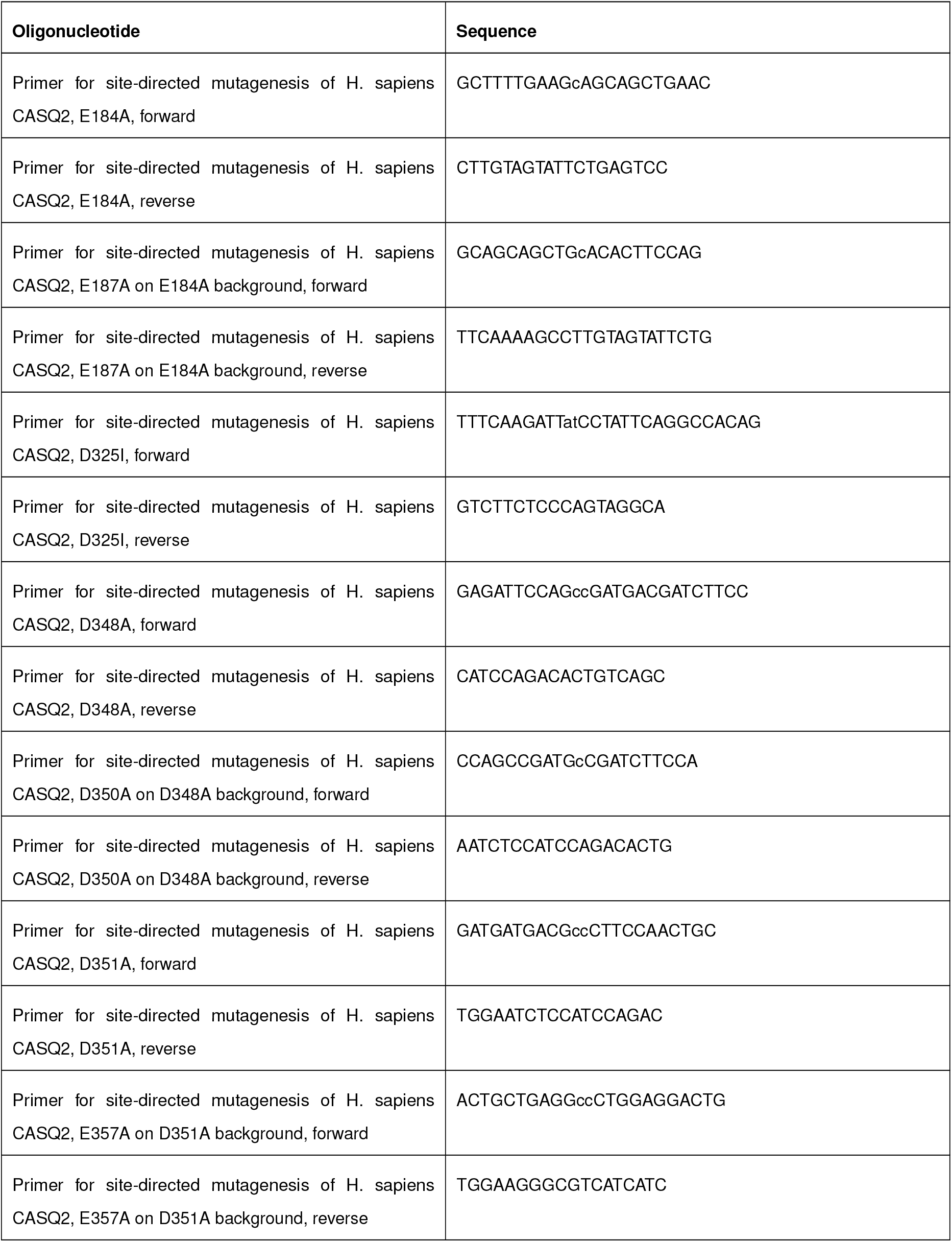
Oligonucleotides Used in This Study

## Notes

#### Summary of Updates

Shortened title and abstract; minor textual changes; cosmetic changes to figures.

https://github.com/errontitus/casq2-structure-function

